# Two dynamically distinct circuits driving inhibition in sensory thalamus

**DOI:** 10.1101/2020.04.16.044487

**Authors:** Rosa I. Martinez-Garcia, Bettina Voelcker, Julia B. Zaltsman, Saundra L. Patrick, Tanya R. Stevens, Barry W. Connors, Scott J. Cruikshank

## Abstract

Most sensory information destined for the neocortex is relayed through the thalamus, where considerable transformation occurs^1,2^. One powerful means of transformation involves interactions between excitatory thalamocortical neurons that carry data to cortex and inhibitory neurons of the thalamic reticular nucleus (TRN) that regulate flow of those data^3-6^. Despite enduring recognition of its importance^7-9^, understanding of TRN cell types, their organization, and their functional properties has lagged that of the thalamocortical systems they control.

Here we address this, investigating somatosensory and visual circuits of the TRN. In the somatosensory TRN we observed two groups of genetically defined neurons that are topographically segregated, physiologically distinct, and connect reciprocally with independent thalamocortical nuclei via dynamically divergent synapses. Calbindin-expressing cells, located in the central core, connect with the ventral posterior nucleus (VP), the primary somatosensory thalamocortical relay. In contrast, somatostatin-expressing cells, residing along the surrounding edges of TRN, synapse with the posterior medial thalamic nucleus (POM), a higher-order structure that carries both top-down and bottom-up information^10-12^. The two TRN cell groups process their inputs in pathway-specific ways. Synapses from VP to central TRN cells transmit rapid excitatory currents that depress deeply during repetitive activity, driving phasic spike output. Synapses from POM to edge TRN cells evoke slower, less depressing excitatory currents that drive more persistent spiking. Differences in intrinsic physiology of TRN cell types, including state-dependent bursting, contribute to these output dynamics. Thus, processing specializations of two somatosensory TRN subcircuits appear to be tuned to the signals they carry—a primary central subcircuit to discrete sensory events, and a higher-order edge subcircuit to temporally distributed signals integrated from multiple sources. The structure and function of visual TRN subcircuits closely resemble those of the somatosensory TRN. These results provide fundamental insights about how subnetworks of TRN neurons may differentially process distinct classes of thalamic information.

## Two types of neurons in the somatosensory TRN

We first explored the cellular composition of the TRN, characterizing expression of parvalbumin (PV), calbindin (CB), and somatostatin (SOM)—three markers found to be useful in differentiating functionally distinct neural types in the neocortex and elsewhere^13^. Brain sections spanning the somatosensory sector of the TRN^4,7,8^ were prepared from SOM-Cre mice crossed with Cre-dependent tdTomato (tdT) reporters, then stained immunohistochemically for CB and PV (Fig. 1a). Nearly all somatosensory TRN cells expressed PV^1,14^, whereas only subsets expressed SOM-tdT or CB (∼64% and ∼48%, respectively; Fig. 1c; Extended Data Fig. 1).

**Figure 1.**
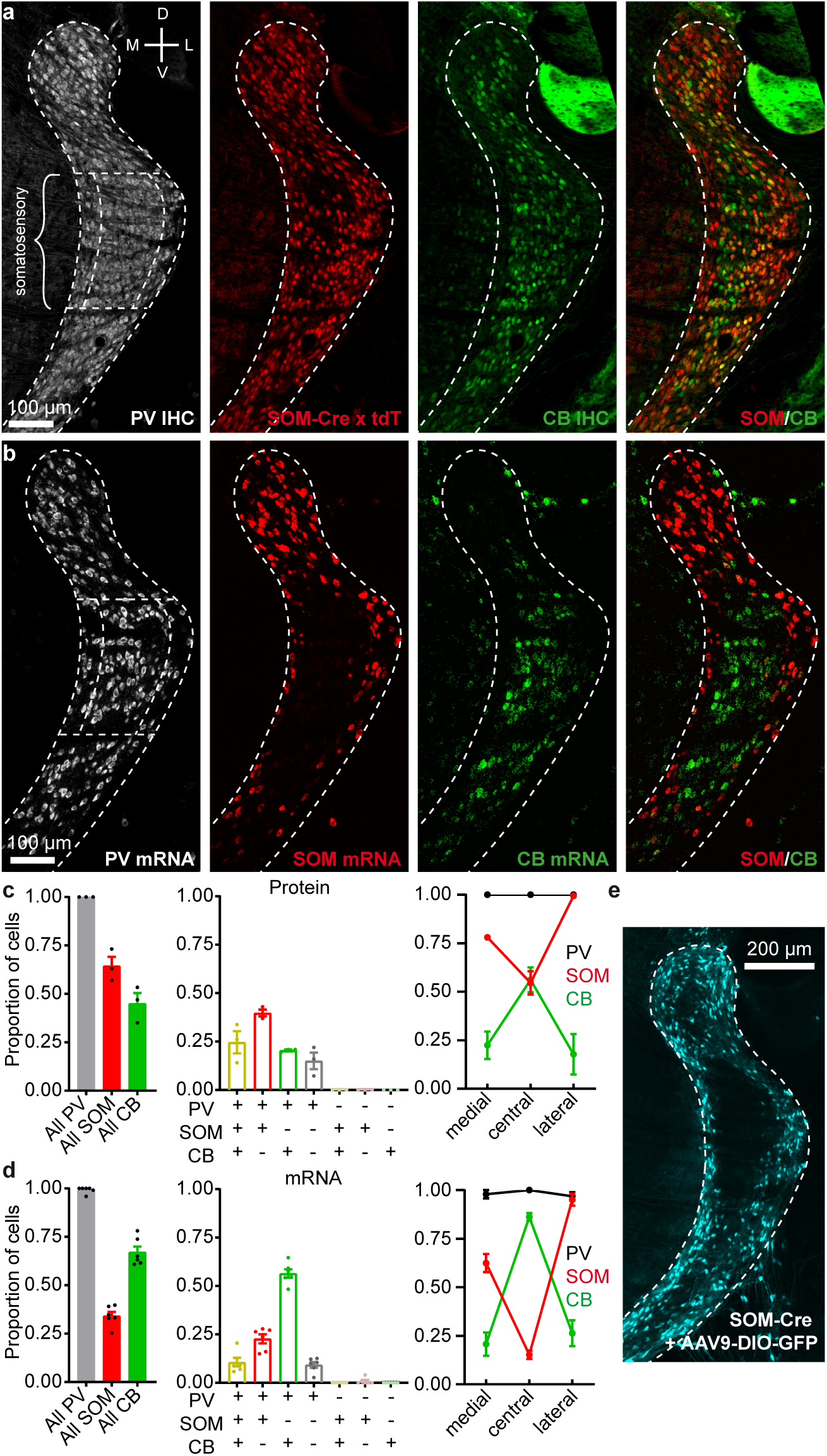
Somatosensory TRN is composed of neurochemically distinct neurons located in separate zones. **a**, Confocal images of a 40 µm section centered on somatosensory TRN, from a SOM-Cre x tdTomato mouse. Immunohistochemical (IHC) staining for parvalbumin (PV) and calbindin (CB). Somatostatin-Cre cells were genetically labeled (SOM-Cre x tdT). TRN outlines based on PV labeling. **b**, Confocal images of an 18 µm section from a WT mouse showing FISH for *Pvalb, Sst, and Calb1* (PV, SOM, CB). **c**, Left, proportion of cells expressing PV, SOM-Cre x tdT, or CB in somatosensory TRN (bracketed sector, above). Middle, proportions of cells expressing combinations of markers (n=1075 cells, 3 sections, 3 mice). Right, proportions for each marker in 3 zones of somatosensory TRN. 187 cells were in the medial zone (medial 20%), 746 in the central zone (central 60%), and 142 in the lateral zone (lateral 20%). PV images show zones. Fractional SOM-Cre x tdT cell densities were higher in the edge than central zones, whereas CB cell densities were higher centrally (all *P* < 0.001, chi-square, Yates’ correction). **d**, Same as (c) but for mRNA (n=593 cells, 6 sections, 6 mice). Right, 98 cells were in the medial zone, 412 in the central, and 83 in the lateral. Again, SOM and CB cells were differentially distributed across zones (all P < 0.001, chi-square, Yates’ correction). **e**, Confocal image of a 90 µm slice from a P23 SOM-Cre mouse. AAV9-DIO-GFP was injected in TRN at P14 to assay SOM-Cre expression. GFP cells (pseudocolored cyan) were largely absent in the central zone (Replicated in 12 mice). Data expressed as mean ± SEM.

SOM-tdT and CB cells had complementary distributions across the somatosensory TRN. SOM-tdT cells were at highest densities near the medial and lateral edges of the sector, whereas CB cells were concentrated near the center and virtually absent along the edges (Fig. 1a; Extended Data Fig. 1). Quantitative comparisons between the medial 20%, lateral 20% and central 60% of the somatosensory TRN confirmed that proportions of cells expressing SOM-tdT were higher in the edge zones than the central zone (p < 0.0001), while the reverse was true for CB (p < 0.0001) (Fig. 1c).

To test this organization further, we used *in situ* hybridization to assay SOM, CB, and PV mRNA expression. The highest SOM cell densities were again in the medial and lateral edge zones, CB cells were clustered centrally, and nearly all cells were PV positive (Fig. 1b, d; Extended Data Fig. 2). Interestingly, the edge/central segregation of SOM/CB cells was more salient in the mRNA assay, mainly due to decreased proportions of SOM cells in the central zone (only 15.3% of central cells expressed SOM mRNA whereas 54.7% expressed SOM-Cre x tdT; Fig. 1a-d).

The near absence of SOM mRNA in the central zone suggests that most neurons located there may not actually express SOM protein in mature animals, and that central cell expression of tdT in SOM-Cre x tdT mice could result from genetic recombination early in development, and persistent tdT production thereafter^15^. To test for mature SOM expression, we initially tried immunohistochemistry but were unable to find SOM antibodies adequate for TRN (not shown). As an alternative, we assayed Cre expression in mature SOM-Cre mice, injecting adeno-associated virus driving Cre-dependent GFP in TRN. Cre expression in somatosensory TRN of these mice was almost entirely restricted to the edge zones, consistent with the SOM mRNA pattern (Fig. 1b, d-e, Extended Data Fig. 3). Together, the results indicate that somatosensory TRN is composed of neurochemically distinct cell types segregated into separate zones: a core central zone is composed mostly of CB-expressing neurons, and it is flanked by edge zones of SOM-expressing neurons.

## Primary and higher-order TRN subcircuits

First-order VP and higher-order POM thalamocortical (TC) nuclei transmit distinct information to different targets in the neocortex and send collaterals to the TRN, the latter leading to both open- and closed-loop thalamic inhibition^4-6,8^. Clarifying the organization of these circuits, including how first-order and higher-order TC nuclei synapse with subtypes of TRN neurons^16^, is essential to understanding thalamic information processing^7^. To this end, we selectively expressed ChR2-eYFP in VP or POM, then characterized their inputs to TRN (Fig. 2). Strikingly, their projections segregated topographically in close alignment with the observed patterns of TRN cell types. VP axons terminated in the CB-rich central zone of somatosensory TRN, whereas POM axons terminated along the SOM-dense medial and lateral edges (Fig. 2a-b; Extended Data Fig. 4a, c).

**Figure 2.**
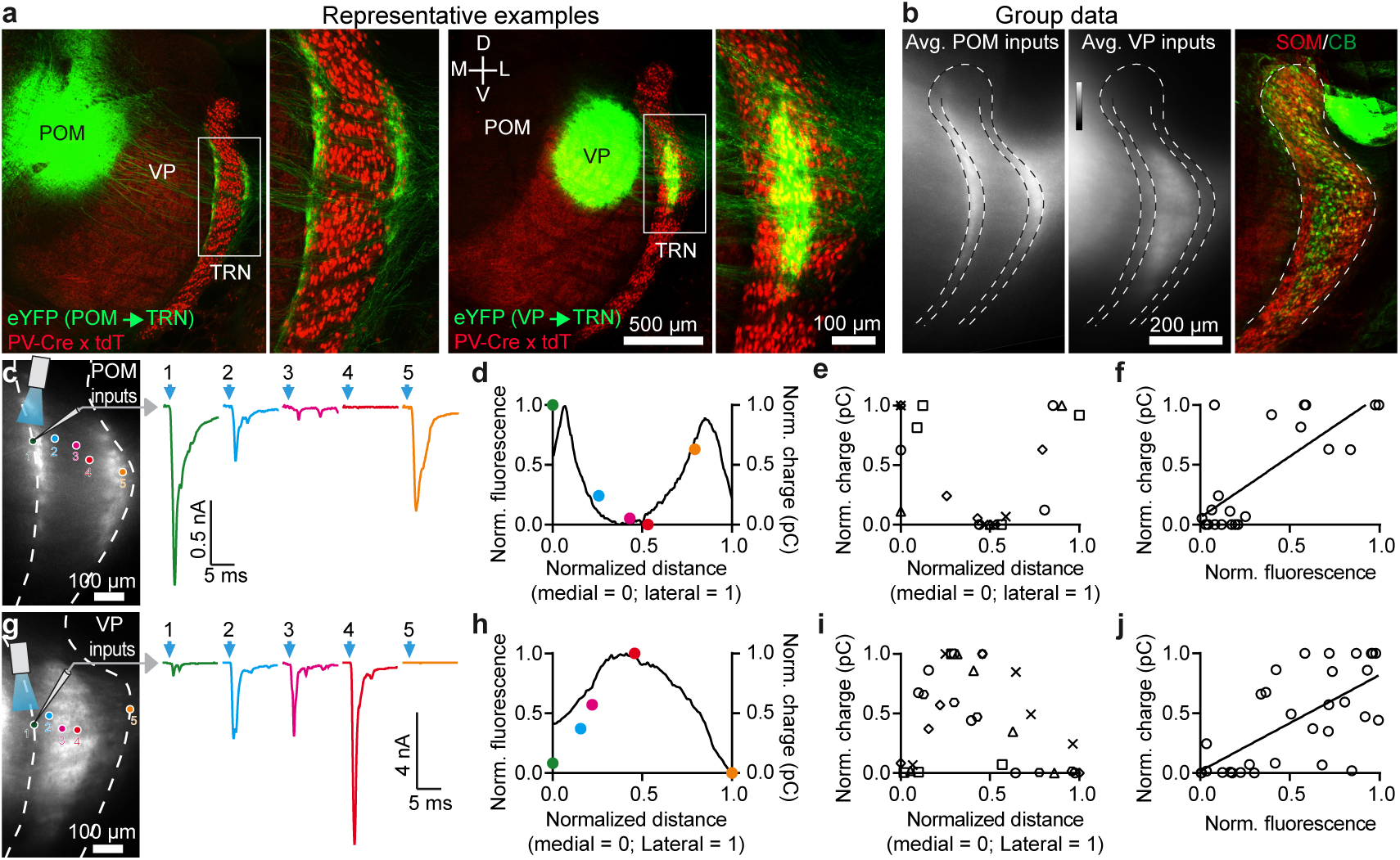
VP, a first-order thalamic nucleus, targets central somatosensory TRN, whereas POM, a higher-order nucleus, targets the TRN edges. **a**, Confocal images illustrating the distinct projections from POM and VP to TRN. AAV2-ChR2-eYFP was injected in POM (left) or VP (right) and projections to TRN characterized. **b**, Group data showing average thalamic projections to TRN. Viral anterograde tracing as in (a). POM targeted the edges of somatosensory TRN (n=9 mice). VP targeted the central zone (n=8 mice). Central/edge boundaries drawn at 20% and 80% of medial-lateral distance across TRN. Right, average SOM/CB immunohistochemical profiles (n=5 mice) aligned to same reference TRN as anterograde projection maps. **c**, Synaptic responses to POM input from cells across the TRN, testing how synaptic strength relates to anterograde fluorescence and topographical location. Left, image of live TRN (outlined) with ChR2-eYFP projections from POM. Circles show locations of recorded cells. Right, EPSCs evoked in TRN cells by optical activation of POM axons (−84 mV). Colors match cells in left image and (d). **d**, Normalized fluorescence (black line) and synaptic charge (colored dots) for cells in (c), versus medial-lateral position in TRN. **e**, Group relationship between TRN soma location and evoked synaptic response. Somas close to the TRN edges responded strongly to POM and those in the center did not (for e-f, n=21 cells, 5 slices, 5 mice; each prep has unique symbol). **f**, TRN cells’ synaptic responses to POM input (normalized charge) correlated with fluorescence from POM axons surrounding the cell (r=0.75, p<0.0001, two-tailed Pearson’s correlation). **g-h**, Same as (c-d), except activating VP input. **i**, Group data showing that somas near the TRN center respond more strongly to VP input than those near the edges (for i-j, n=31 cells, 6 slices, 4 mice). **j**, TRN cell’s synaptic responses to VP input correlated with fluorescence from VP axons surrounding the cell (r=0.68, p<0.0001, two-tailed Pearson’s correlation).

Given this stark anatomical segregation of projections, it seemed likely that central and edge TRN cells would be selectively targeted by synapses from VP and POM, respectively. However, the dendrites of TRN neurons might extend into adjacent zones, leading to functional cross-talk among the circuits^1,8,17^. To address this, we mapped excitatory synaptic strengths of POM and VP inputs to cells located across the mediolateral axis of somatosensory TRN. Consistent with the anatomy, synaptic responses to POM inputs were much stronger for TRN edge cells than for central cells, whereas VP inputs evoked strongest responses in central cells (Fig. 2c-e, g-i; Extended Data Fig. 4b, d). Moreover, synaptic strengths for TRN cells correlated with fluorescence intensities of the afferent terminals near their soma (Fig. 2d, f, h, j). Together, our findings show that primary and higher-order somatosensory thalamic inputs to TRN are topographically segregated and align with the neurochemical pattern of TRN cell types: VP projects strongly to CB-expressing central cells, and POM to SOM-expressing edge cells.

## TRN subcircuits are functionally distinct

Primary and higher-order thalamocortical nuclei convey qualitatively different types of information^10,12,18-21^, and we wondered if VP and POM communication with TRN might involve parallel differences in synaptic mechanisms. To address this, we compared dynamic features of the glutamatergic synaptic currents (kinetics and short-term synaptic depression) evoked by photostimulating VP and POM inputs to TRN. VP synaptic currents in central TRN cells were brief and depressed deeply during repetitive activation. Conversely, POM currents in TRN edge cells were longer-lasting and more stable, depressing significantly less (Fig. 3a-c, Extended Data Fig. 5-6).

**Figure 3.**
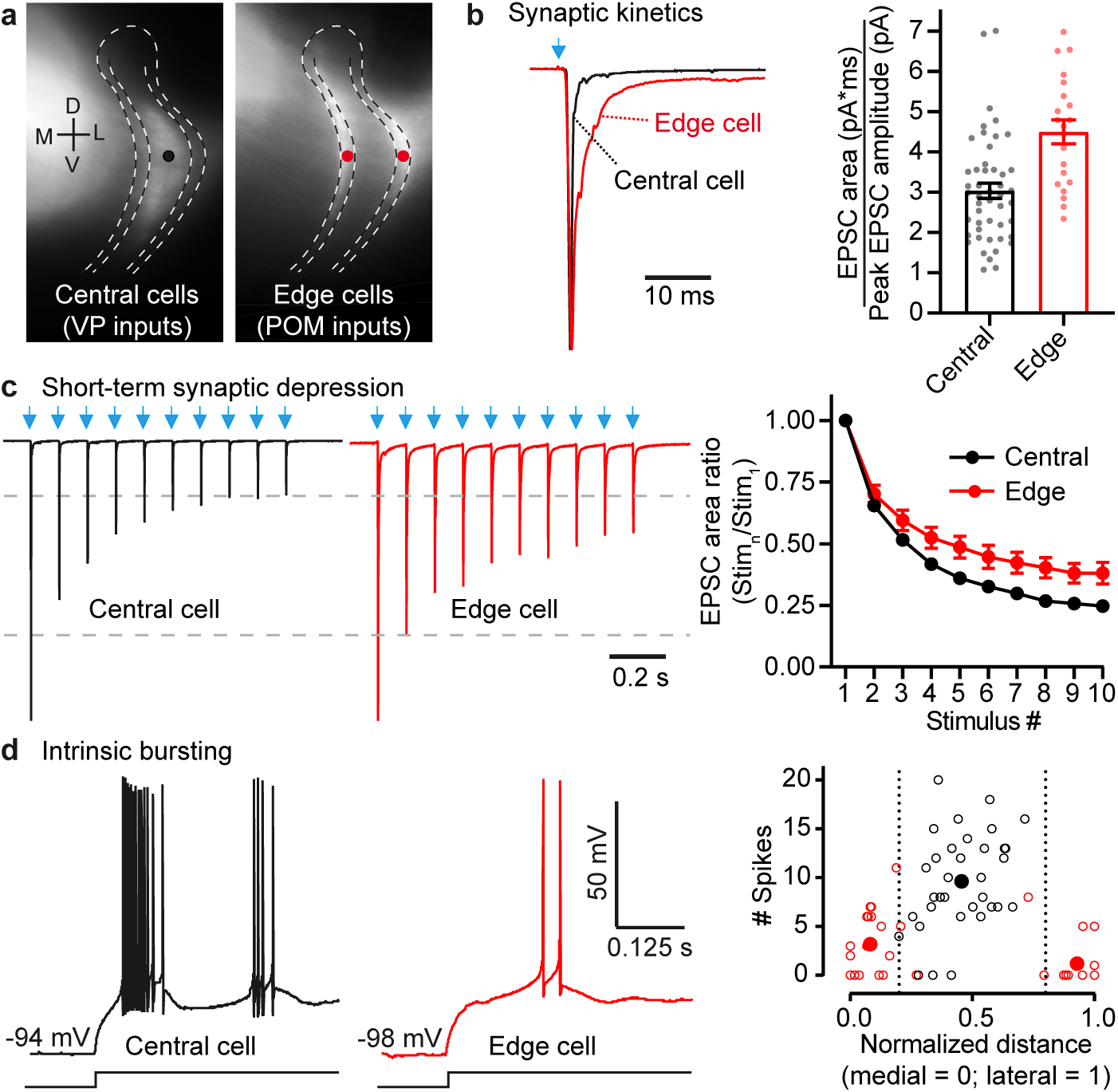
Central and edge TRN cells receive thalamic inputs with different kinetics and synaptic depression, and have distinct bursting tendencies. **a**, Representative locations of TRN cells targeted by VP (black dot) and POM (red dots). **b**, POM-evoked EPSCs of edge cells are more prolonged than VP-evoked EPSCs of central cells. Left, normalized optogenetically evoked example EPSCs for each cell type (stimulus 1 of (c); V_m_ -84 mV). Right, POM-evoked edge cell responses have greater fractional area in later stages of the EPSCs (p<0.0001, unpaired two-tailed t-test; central: 48 cells, 14 mice; edge: 21 cells, 14 mice). **c**, Short-term synaptic depression. Left, representative EPSCs from central cells (VP inputs) and edge cells (POM inputs), normalized to response peaks (∼2 nA) (10 Hz LED trains). Right, short-term depression was greater for central than edge cell groups (p < 0.0001, ANOVA stim. 2-10; all p < 0.005 from stimulus 5-10, two-tailed Bonferroni’s t-test; same cells as (b)). **d**, Left, intrinsic bursting of example central and edge cells. Offset bursts were triggered by injecting negative current to reach ∼-95 mV, followed by abrupt current removal. Right, number of spikes per burst as a function of position in TRN (open circles: individual cells; filled circles: zone averages; 34 central cells, 11 mice; 22 medial cells, 10 mice; 16 lateral cells, 10 mice). Edge cells (red) discharged fewer spikes/burst than central cells (black) (unpaired two-tailed t-test, p<0.0001). Data expressed as mean ± SEM.

To understand how these dynamically distinct inputs to TRN neurons might be integrated postsynaptically, we assessed the cells’ intrinsic physiological characteristics. Central and edge neurons differed across a range of passive and active membrane properties^17^. Edge cells had higher resistances, lower capacitances, and smaller somata than central cells. Action potential kinetics, afterpotentials, and threshold currents also differed (Extended Data Fig. 7; Supplementary Information 1). One striking intrinsic distinction among TRN cell types, which could powerfully influence responses to synaptic input during certain behavioral states^1,2,5,6,22-26^, was the much greater tendency of central cells to fire spikes in high frequency bursts (Fig. 3d). The bursting differences were consistent with stronger T-type (low threshold) calcium currents in central cells^17,27-30^. Thus, nearly all central cells fired “offset bursts” following release from hyperpolarizing stimuli (∼10 spikes/burst). In contrast, under matched stimulus conditions, most edge cells either failed to burst or fired weak bursts (∼2 spikes/burst; Fig. 3d; Supplementary Information 1). Neither cell type exhibited bursting when excited from a more depolarized steady-state (∼-74 mV; Extended Data Fig. 7).

We next examined how the observed pathway-specific synaptic and intrinsic properties combine to control spiking responses of TRN cells to their excitatory thalamic inputs. The brief and depressing synaptic inputs from VP to central TRN cells, together with the propensity of central cells to burst, suggests they might respond phasically – initially strong but quickly decreasing with repeated activation. In contrast, the kinetically slow and more stable inputs from POM to the less bursty edge cells predict initially weaker but more sustained spiking.

First, we generated simulated synaptic currents that matched the EPSCs previously recorded from central and edge TRN cells in response to their respective inputs. We then characterized TRN spike responses elicited by intracellular injection of these currents, delivered while the TRN cells were at their resting potentials (∼-84 mV). As predicted, central cells responded to simulated VP inputs with initial bursts that sharply depressed. In contrast, spiking responses of edge cells to simulated POM inputs were initially much weaker but persisted more during repetitive activation (Fig. 4a-b).

**Figure 4.**
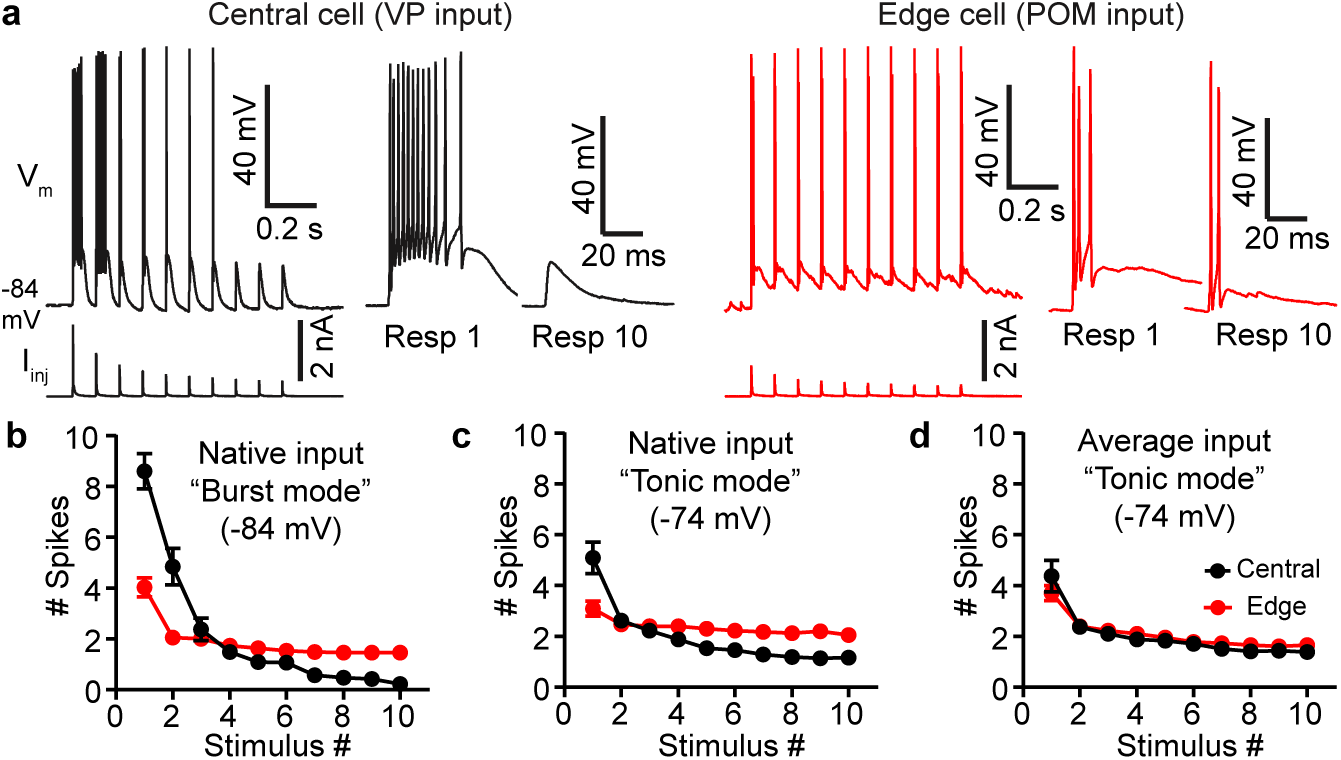
Thalamus-evoked spiking in central TRN cells is transient, whereas edge cell responses are more sustained. **a**, Simulated synaptic currents, modeled as average recorded EPSCs evoked by activation of VP or POM axons (Fig 3; Methods), were applied to central and edge TRN cells. Left (black), response of a central cell (V_m_) to simulated VP synaptic currents (I_inj_). The first stimulus evoked a 12-spike burst, but responses depressed to zero spikes by stimulus 8. Right (red), response of an edge cell to simulated POM synaptic currents. The first stimulus evoked 3 spikes, and spiking persisted during repetitive stimulation. **b**, Mean spike outputs of central cells to simulated VP currents, and edge cells to simulated POM currents (i.e. to their native synaptic inputs). TRN cells were held in burst mode (−84 mV). Central cell responses were initially strong and depressed sharply, whereas edge responses were more stable (spike counts from stimulus 1 to stimulus 10 declined by 97% for central cells and 63% for edge cells; p<0.0001, unpaired two-tailed t-test). For (b)-(d): 13 central cells, 11 edge cells, 8 mice. **c**, Same as (b) except cells held at -74 mV to reduce intrinsic bursting. Central responses became more edge-like, yet significant differences remained (spike counts from stimulus 1 to stimulus 10 declined by 77% for central and 33% for edge cells; p<0.0003, unpaired two-tailed t-test). **d**, Spike responses of central and edge cells (−74 mV) to identical simulated synaptic currents (average of VP and POM EPSCs). Central and edge cell responses did not differ significantly (p=0.19, unpaired two-tailed t-test). Values are mean ± SEM.

Did differences in intrinsic bursting (Fig. 3d) contribute to these striking differences in synaptically evoked spiking? To address this, we depolarized the TRN cells to -74 mV to partially inactivate T-type calcium channels and reduce intrinsic bursting, then recharacterized responses. Central responses became far less phasic and were more persistent, whereas edge cells were barely affected. Initial spiking was reduced 41% in central cells but only 23% in edge cells, and responses to later stimuli in the trains were enhanced more for central cells (Fig. 4c). These results indicate that intrinsic bursting in central TRN cells has a powerful role in their responses to excitatory inputs when that input arrives during relatively hyperpolarized states (e.g., during sleep or periods of strong inhibition). The far smaller effects of polarization on edge cells indicates less influence of T-type calcium bursting and, importantly, weaker modulation by the kinds of membrane potential shifts thought to occur during behavioral state transitions^1,6,22-26,31^.

Finally, we wondered whether the dynamic features of VP synaptic inputs to central cells (faster, more depressing; Fig. 3a-c) also contributed to their phasic spike outputs. For this we generated simulated synaptic currents that were the average, in terms of EPSC kinetics and short-term depression, of the VP → central and POM → edge synaptic currents. We then tested the effects on evoked spiking, with steady-state potentials set to -74 mV to minimize bursting. Remarkably, responses of the two cell types were virtually identical when triggered by the averaged synaptic input; central responses became more sustained and edge responses slightly more phasic (Fig. 4c-d). Together, these results indicate that central and edge cells differentially transform their native excitatory thalamic inputs into distinct spiking outputs through differences in both dynamics of their synaptic inputs and their intrinsic burstiness.

## TRN inhibitory outputs are subcircuit-specific

To better understand the consequences of the distinct output from the TRN subcircuits, we examined their projections and the inhibitory feedback they produced. Interestingly, the two TRN cell types predominantly inhibited the thalamocortical nuclei that drive them. That is, CB-expressing TRN cells projected to and inhibited VPM neurons, whereas SOM-expressing edge cells bypassed VPM and instead inhibited POM (Extended Data Fig. 8). Thus, the primary and higher order segregation of somatosensory reticulo-thalamic subcircuits appears to be largely reciprocal.^8,16,32^

## Visual & somatosensory TRN have similar subcircuits

To test whether other sensory systems might share the salient structural-functional organization described above, we examined the visual TRN and its associated thalamocortical nuclei—the primary dorsal lateral geniculate and the higher-order lateral posterior nucleus (pulvinar). Remarkably, the organization of the visual TRN, in terms of its synaptic input patterns and intrinsic physiological properties of its neurons, was strikingly similar to that of the somatosensory system (Extended Data Fig. 9). This suggests that a primary central core, flanked by higher-order edge neurons, may be a widespread TRN motif.

## Discussion

Our results imply that sensory regions of TRN have two discrete subcircuits distinguished by their structure and function (Extended Data Fig. 10). Structurally, two types of TRN neurons are segregated into central and edge zones and receive inputs from different thalamocortical nuclei. Functionally, the subcircuits have distinct dynamics determined by the intrinsic physiology of their respective neurons and properties of their excitatory thalamic synapses. The subcircuits appear to be tuned to temporal characteristics of the signals they process—transient sensory signals in the primary systems and more temporally distributed signals in the higher-order ones^10,11,18,21^.

A longstanding hypothesis is that the TRN serves as a gatekeeper of information flow, permitting distinct thalamocortical circuits to regulate one another, and enabling functions such as selective attention^1,5,9,33^. It has been proposed that inhibitory cross-talk between thalamic circuits^32,34^ may underlie such regulation^8,35,36^. However, the sharp and reciprocal segregation of subcircuits we observed suggests that intrathalamic cross-talk may play a minor role. Instead, cross-system regulation in the thalamus could be mediated by other means, including descending cortical control of reticulo-thalamic subcircuits^2,37-39^, the nature of which is only beginning to emerge^40^.

The distinct kinetics of the parallel TRN subsystems suggests that arriving signals will be filtered through circuit-specific temporal tuning mechanisms. For example, it is generally thought that TRN neurons undergo shifts from bursting to tonic spiking mode as animals transition between behavioral states of quiescence and aroused wakefulness^1,22,24,26^, due to activity-dependent changes in membrane potentials and neuromodulatory tone^2,6,23,25,28,41^. Our results imply that such state transitions alter spike mode in primary/central TRN cells much more strongly than in higher-order/edge cells^27,31^. This, and the other kinetic differences between subcircuits we describe, likely have profound effects for thalamic processing of afferent signals^5^, both ascending and descending^38^.

Previous studies of several species and sensory systems have suggested separate TRN laminae for primary and higher-order connections^1,7,8^, generally with just a single higher-order layer^16,32,42,43^. In contrast, in the somatosensory TRN of the mouse, we observed a primary core of CB cells surrounded by a higher-order shell-like zone of SOM cells. In at least one respect, this organization is reminiscent of the visual TRN of the primate galago; both systems have SOM-expressing zones that receive inputs from higher-order thalamus, and non-SOM zones receiving primary inputs^43^. Thus the association between SOM neurons of TRN and higher-order processing may be a conserved circuit feature^44^; this, together with the association between CB expression and first-order processing, are especially exciting results because they enable powerful new strategies for probing behavioral and perceptual functions of these distinct TRN circuits^45^.

## Endnotes

## Acknowledgements

We thank Zhanyan Fu, Guoping Feng, and their associates investigating the TRN, for cooperative and stimulating interactions surrounding this project. We also thank Shane Crandall, Omar Ahmed, Mark Zervas, Diane Lipscombe, Chinfei Chen, Gabriela Manzano, Saba Baskoylu, Frederic Pouille, Brian Theyel, and Frank (Scott) Susi for helpful discussions. This work was supported by R01 NS100016, P20 GM103645, NSF 1738633, NSFGRFP 1058262, NSF 1632738.

## Author contributions

R.I.M-G. designed and conducted the experiments, analyzed the results, and wrote the paper. B.V. designed and conducted the experiments, analyzed the results and wrote the paper. J.B.Z. conducted experiments and analyzed the results. S.L.P. and T.R.S. conducted experiments. B.W.C. designed experiments and wrote the paper. S.J.C. designed and conducted the experiments, analyzed the results, and wrote the paper.

## Competing Interests

Authors declare no competing interests.

**Correspondence and requests for materials** should be addressed to S.J.C.

## Data and Materials Availability

Data will be made available by the corresponding author upon request.

---

**Extended Data Figure 1.**
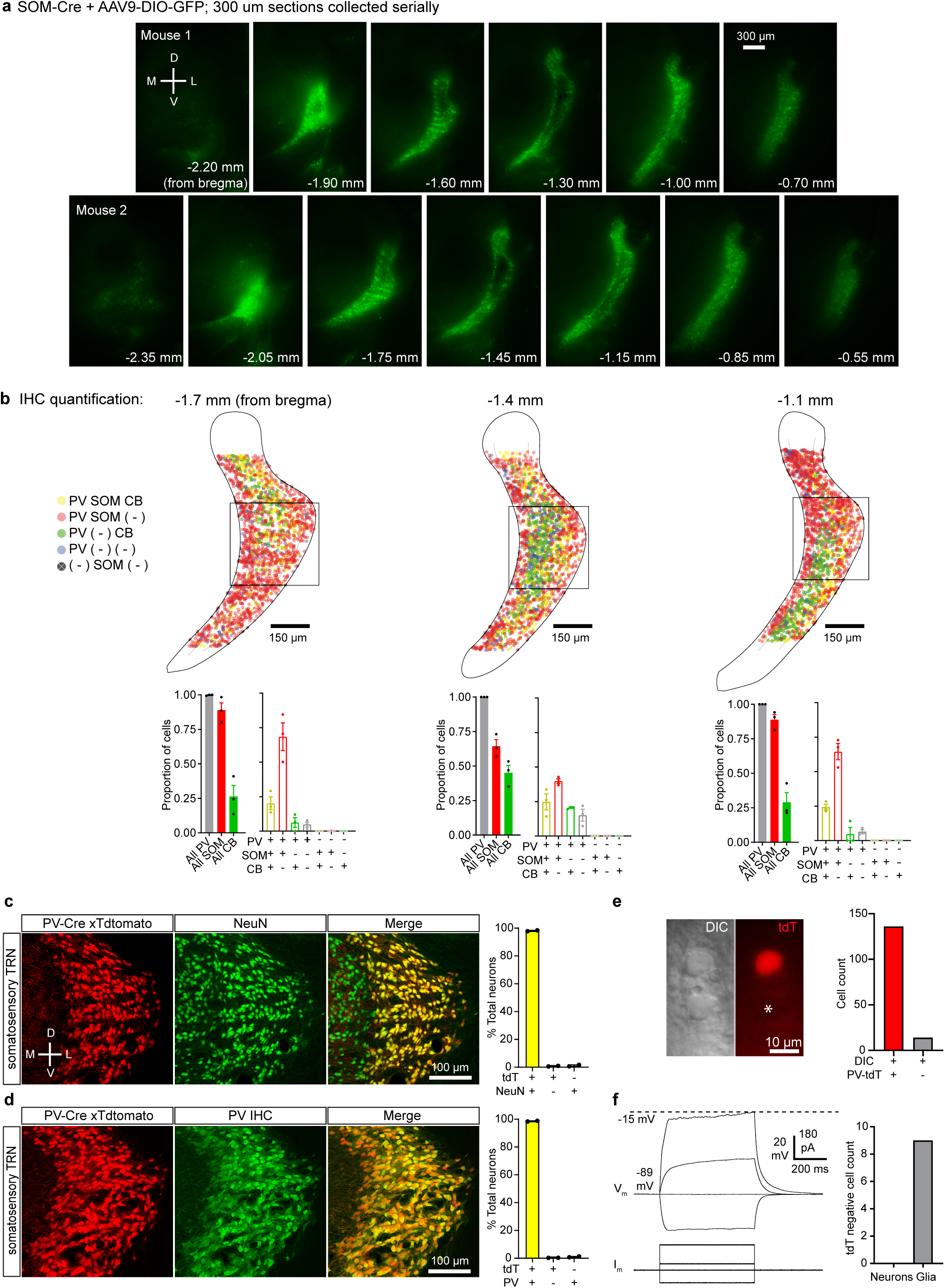
Somatostatin cells form a shell around a calbindin-rich central core of somatosensory TRN; nearly all neurons in somatosensory TRN express parvalbumin. **a**, Epifluorescence images showing serial sections of live tissue expressing AAV9-DIO-GFP (9 days of expression) through the TRN of two P23 SOM-Cre mice. Anterior-posterior (AP) positions of the sections relative to bregma are indicated. Notice how the SOM-expressing cells form a shell-like structure surrounding the central core of somatosensory TRN (centered ∼ -1.4 mm from bregma) (experiment replicated 12 times). **b**, Top, dot plots showing the locations of the genetically or immunohistochemically labeled TRN cell types for sections at different AP distances from bregma. Bottom, proportion of cells expressing PV, CB, SOM-Cre x tdT, and combinations of the markers. Quantification was restricted to the boxed regions shown in the top panels (cell counts: left = 947, middle = 1075, right = 843; 3 sections from 3 mice). The proportion of CB cells was highest in the middle section of somatosensory TRN (−1.4 mm), whereas the proportion of SOM-Cre x tdT cells was lowest in the middle section and higher in the more anterior and posterior sections (P < 0.001, chi-square test with Yates’ correction). **c**, Left, confocal images of the somatosensory sector of TRN in a PV-Cre x tdT mouse (red) stained for NeuN (green). Right, quantification of expression. In PV-Cre x tdT mice, 98.09% of TRN neurons were double positive for tdT and NeuN, 0.73% were only positive for tdT, and 1.18% were only positive for NeuN (1218 neurons, 2 sections, 1 mouse). **d**, Same as (c) but tissue was stained immunohistochemically for parvalbumin. In PV-Cre x tdT mice, 98.66% of TRN neurons were double-positive for tdT and PV, 0.34% were only positive for tdT and 0.34% were only positive for PV (1177 neurons, 2 sections, 1 mouse). **e**, DIC-IR (left) and tdT fluorescence (right) images of TRN in a live slice from a PV-Cre x tdT mouse. Asterisk indicates a soma that did not express tdT. During such live imaging, tdT-negative somata were relatively rare in TRN (>80% of somata visualized in DIC-IR expressed tdT; n=5 slices, 3 mice; right panel). **f**, All tdT-negative cells targeted for recording (n = 8/8) had physiological properties of glia. Left, TRN glial cells had hyperpolarized resting potentials (typically < -85 mV) and no action potentials even during large depolarizations. Data in (b) expressed as mean ± SEM.

**Extended Data Figure 2.**
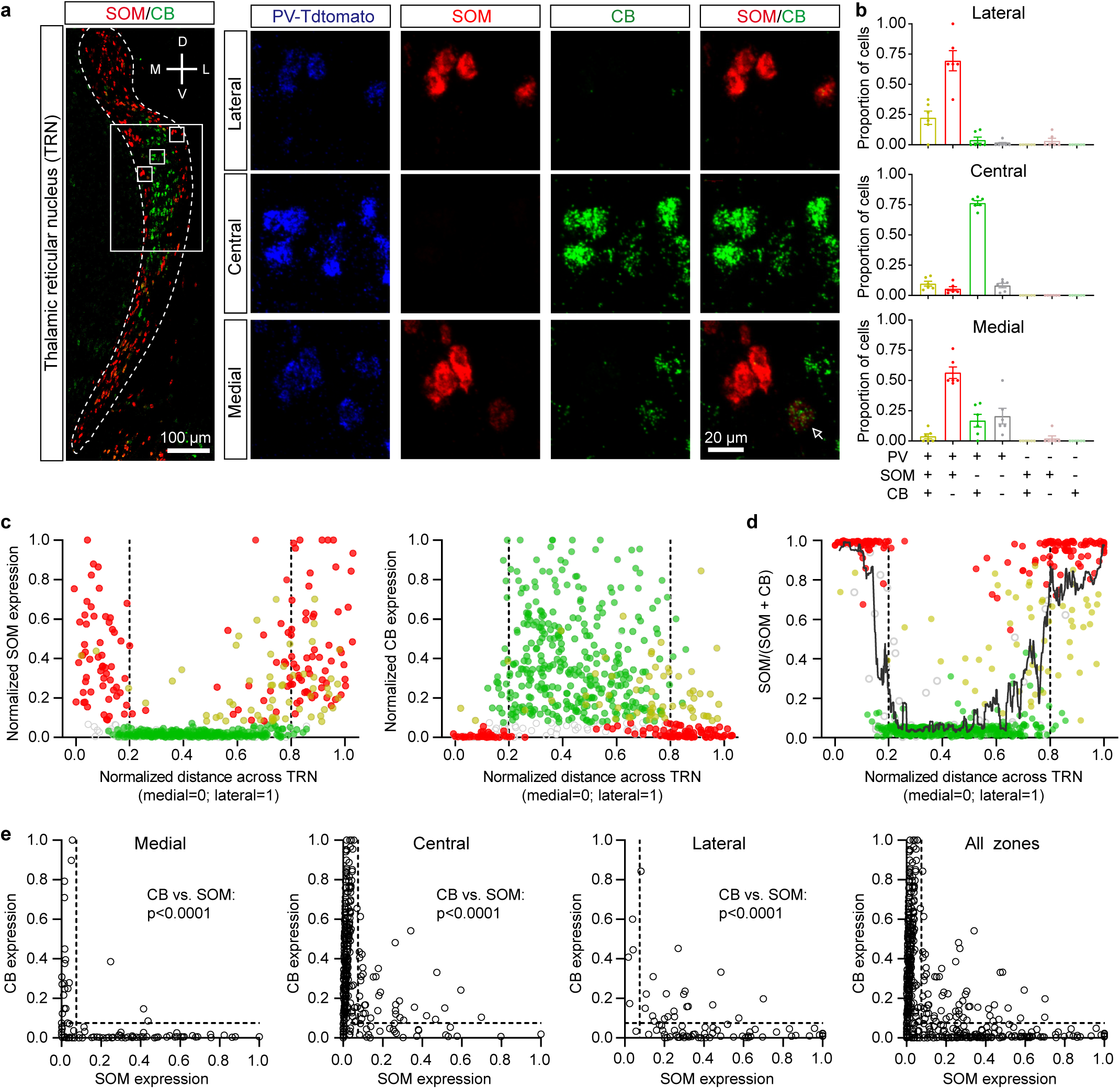
Cells expressing SOM and CB mRNA are topographically segregated in somatosensory TRN. **a**, Left, confocal image overlay of SOM and CB mRNA expression through the TRN of a PV-Cre x tdT mouse. In somatosensory TRN (boxed region), SOM expressing cells are preferentially localized in the medial and lateral edge zones, whereas CB cells are generally in the central zone. Insets show mRNA expression patterns of cells in the lateral edge (top row), central (middle row), and medial edge (bottom row) zones. Although most cells predominantly expressed either SOM (red) or CB (green) mRNA, some (∼10%) expressed both (arrow). **b**, Proportion of cells expressing various combinations of markers for each zone (for b-e, 593 cells, 6 sections, 6 mice; average of 98.9 ± 8.5 cells/section). 99.1 ± 0.2% of cells expressed PV. Most cells in the medial and lateral edge zones expressed only SOM and PV, but not CB (∼64**%**). In contrast, most central cells expressed only CB and PV, but not SOM (∼76**%**). **c**, Normalized SOM (left) or CB (right) mRNA expression as a function of medial-lateral position in the TRN (0= most medial, 1= most lateral). Cells expressing SOM but not CB (red) were concentrated near the edges (mean SOM expression for edge vs central: 0.325 ± .020 vs. 0.052 ± 0.006, p<0.0001, unpaired two-tailed t-test). Cells in the center generally had high levels of CB but not SOM (mean CB expression for central vs edge: 0.326 ± 0.012 vs. 0.086 ± 0.012, p<0.0001, unpaired two-tailed t-test). ∼10% of somatosensory TRN cells expressed both SOM and CB (yellow) and clustered in the lateral half of the TRN, with their focus being near the boundary between the central and lateral zones (average normalized medial-lateral position = 0.71 ± 0.02). Compared with the cells expressing SOM or CB “alone”, the dual SOM/CB cells had lower expression of both SOM (SOM/CB cells: 0.25 ± 0.02 vs SOM only: 0.43 ± 0.02, p<0.0001, unpaired two-tailed t-test) and CB mRNA (SOM/CB cells: 0.23 ± 0.15 vs CB only: 0.39 ± 0.01, p<0.0001, unpaired two-tailed t-test). Together this suggests they may represent a transitional population. About 9% of somatosensory TRN cells expressed PV only (gray outline) and were dispersed fairly evenly across the sector (mean medial-lateral position: 0.38 ± 0.03). **d**, Relative expression of SOM versus CB (within-cell) as a function of medial-lateral position in the TRN (a value of 1.0 on the y-axis index indicates SOM expression only, a value of 0 indicates CB only, and 0.5 indicates equal expression values for SOM and CB). Most cells (80%) had very high (>0.9) or very low (<0.1) relative expression scores, indicating most cells have strong biases toward a single marker (10:1 ratios or greater). Consistent with Fig. 1 and panels a-c here, SOM-biased cells tended to cluster toward the TRN edges, whereas CB-biased were located centrally (edge vs. central preference scores: 0.76 ± 0.03 vs. 0.14 ± 0.01, p<0.0001, paired two-tailed t-test). A moving average (11-point) is superimposed to illustrate the spatial trends in expression bias. Interestingly, even the dual-expressing SOM/CB cells (yellow) tended to have a topographical preference according to relative SOM vs. CB expression: dual-expressing edge cells tended to have higher SOM:CB preference scores than dual-expressing central cells (dual edge vs central cells: 0.58 ± 0.04 vs. 0.47 ± 0.04, p=0.06, unpaired two-tailed t-test). **e**, CB expression as a function of SOM expression for cells in the medial, central, lateral, and combined zones. P-values shown in the panels are the results of two-tailed paired (within-cell) t-tests comparing CB and SOM expression. Data expressed as mean ± SEM.

**Extended Data Figure 3.**
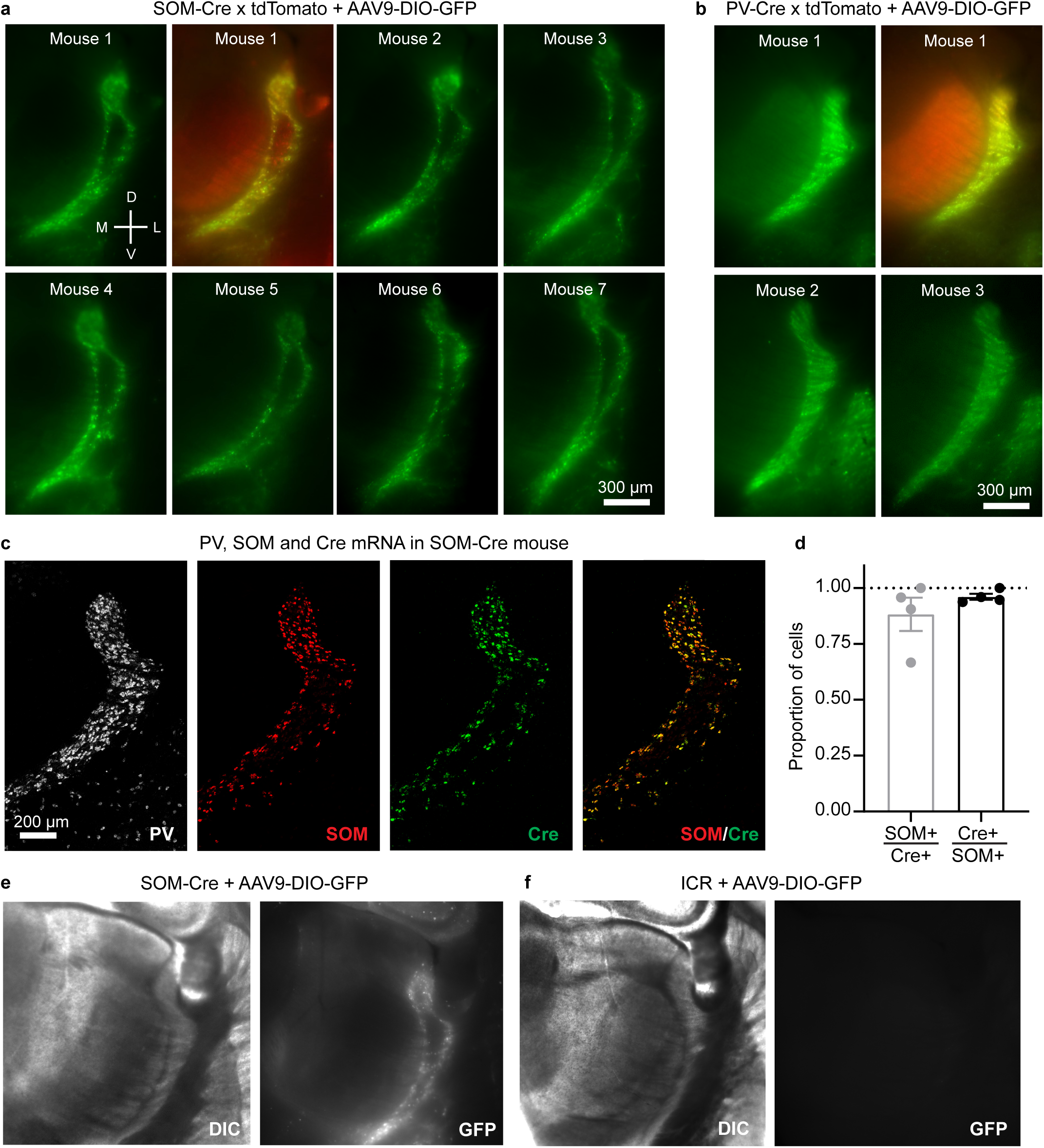
In the somatosensory TRN of postnatal SOM-Cre mice, both Cre expression and viral Cre-dependent protein expression are restricted mainly to SOM cells along the edges. **a**, Epifluorescence images of virally induced, SOM-Cre-dependent GFP expression patterns in TRN (live 300 µm thick slices from 7 mice). Virus carrying a Cre-dependent GFP gene (AAV9-DIO-GFP) was injected throughout the TRNs of SOM-Cre x tdTomato or SOM-Cre x ICR mice. This drove GFP expression (green) in SOM cells that expressed Cre at or after the time of the virus injection (age at injection = P17.1 ± 1.18 days; age at imaging = 26.8 ± 1.19). Mouse 1: images of GFP alone and overlay of GFP and genetically labeled tdTomato cells. Mouse 2-7: GFP alone. Within the somatosensory TRN, expression of SOM-Cre X GFP was restricted to the edges of the nucleus and excluded from the central zone after early development. Experiment replicated 12 times. **b**, Same as in panel (a) except using PV-Cre mice (age at virus injection = P14; age at imaging = P23). Nearly all of the TRN cells expressed both genetically labeled PV-Cre x tdTomato and virally-induced PV-Cre x GFP. In addition to demonstrating ubiquitous postnatal PV-Cre expression across TRN, these results show that the virus has no obvious tropisms for different topographic regions of the TRN. Experiment replicated 4 times. **c**, Confocal images of PV, SOM, and Cre mRNA expression through the TRN of a P27 SOM-Cre heterozygous mouse (18 µm thick section). The overall patterns of SOM and Cre mRNA labeling were nearly identical (Experiment replicated in 5 sections from 4 mice). **d**, SOM and Cre mRNA were generally expressed in the same neurons (total counts: 167 cells, 5 sections, 4 mice; measured in the somatosensory sector of TRN). Of the Cre-expressing cells, 88.3 ± 0.1% expressed SOM mRNA. Of the SOM-expressing cells, 96.1 ± 0.01% expressed Cre mRNA. **e**, AAV9-DIO-eGFP injections into the TRNs of postnatal SOM-Cre mice led to eGFP expression patterns that were very similar to SOM mRNA patterns. Left, DIC and epifluorescence images from a live 300 µm thick section expressing AAV9-DIO-GFP (9 days after virus injection) in a SOM-Cre heterozygous mouse. This experiment was replicated 12 times (see panel a above, Fig. 1g, and Extended Data Fig.1a for additional examples). **f**, Same as (e) but virus was injected into a WT mouse (ICR strain; no Cre expression) (tested in n=2 mice). Matching display settings were utilized for the slices in (c) and (d). The lack of fluorescent signal indicates that the AAV-DIO-GFP virus used here drives GFP expression only in Cre-expressing cells. Data expressed as mean ± SEM.

**Extended Data Figure 4.**
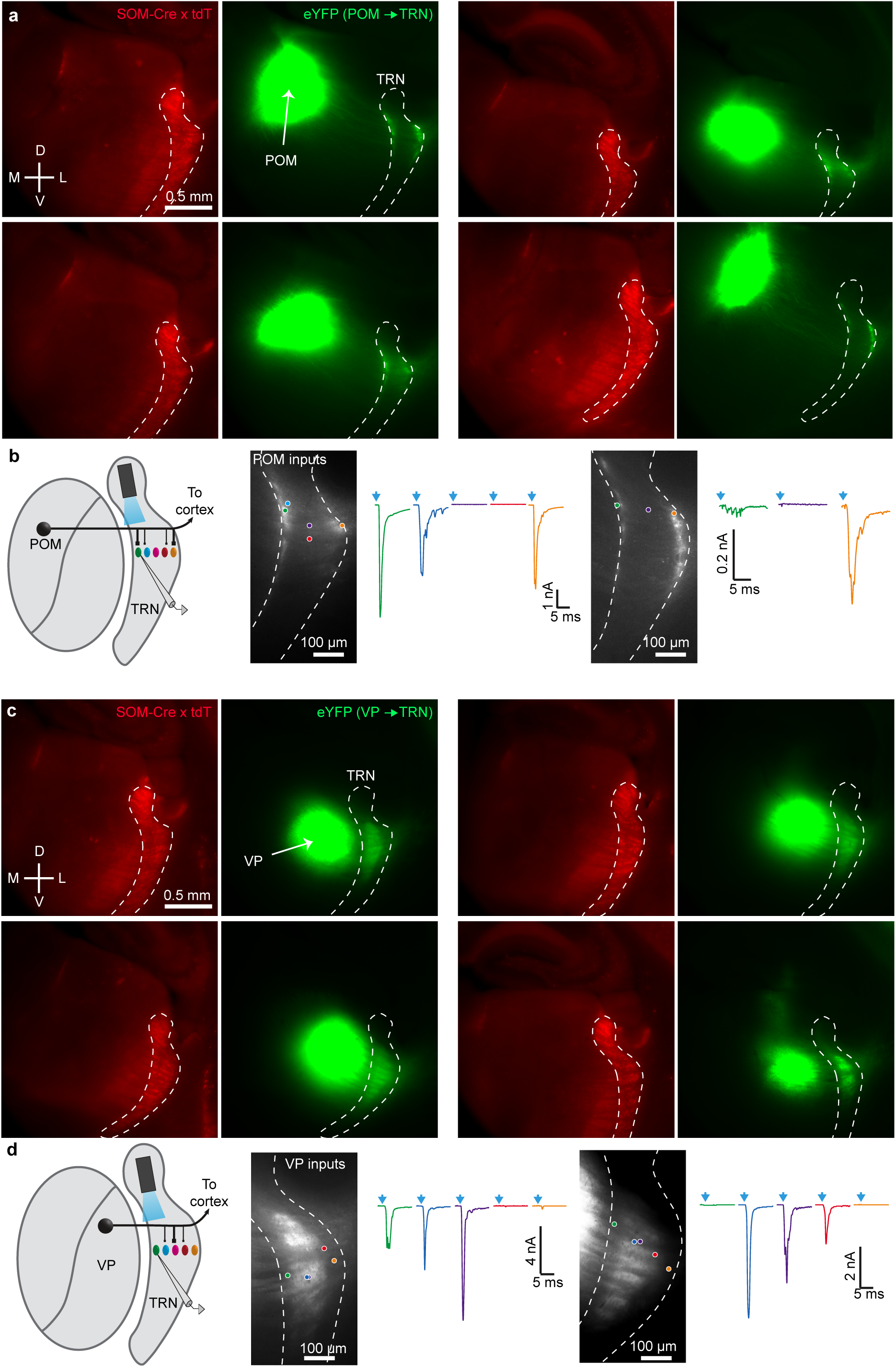
VP afferents target the CB-dense center of somatosensory TRN, whereas POM afferents target the SOM-dense medial and lateral edges. **a**, Four representative examples showing the POM injection sites and anterograde projections to TRN in live 300 µm slices. Red: SOM-Cre x tdT. Green: ChR2-eYFP. Boundaries of the thalamic nuclei (TRN outlined by dashed lines) were determined from SOM-Cre X tdT fluorescence and by brightfield images (not shown). Injection sites were located in POM nuclei, seen here as the large, bright eYFP spots medial to TRN. The smaller bands of eYFP along the medial and lateral edges of the somatosensory TRN are axons and terminals from the labeled POM cells. In addition to the TRN edges, POM also targeted layer 5a (L5a) and L1 of barrel cortex (but not L4; data not shown). Top left image shows the injection site for Fig. 2c-d. Experiment replicated in 18 mice. **b**, Left, schematic showing recording configuration (also used in the experiments of Fig. 2c-f). Right, additional representative examples of synaptic responses of cells across the medial-lateral axis of somatosensory TRN evoked by POM stimulation. TRN cells near the edges of somatosensory TRN received strong input from POM whereas cells located in the center of TRN did not. Experiments replicated in 5 slices, 5 mice. **c**, Similar to a, but for VP afferents. VP afferents clearly targeted the central zone of TRN. VP also projected to L4 of barrel cortex but not L5a or L1 (not shown). Top left image shows the injection site for Fig. 2g-h. Experiments replicated in 25 mice. **d**, Similar to (b), but for VP. Left, schematic showing recording configuration (also used in the experiments of Fig. 2g-j). Right, cells in the center of somatosensory TRN received stronger input from VP than did cells located near the edges of TRN. Experiment replicated in 6 slices, 4 mice.

**Extended Data Figure 5.**
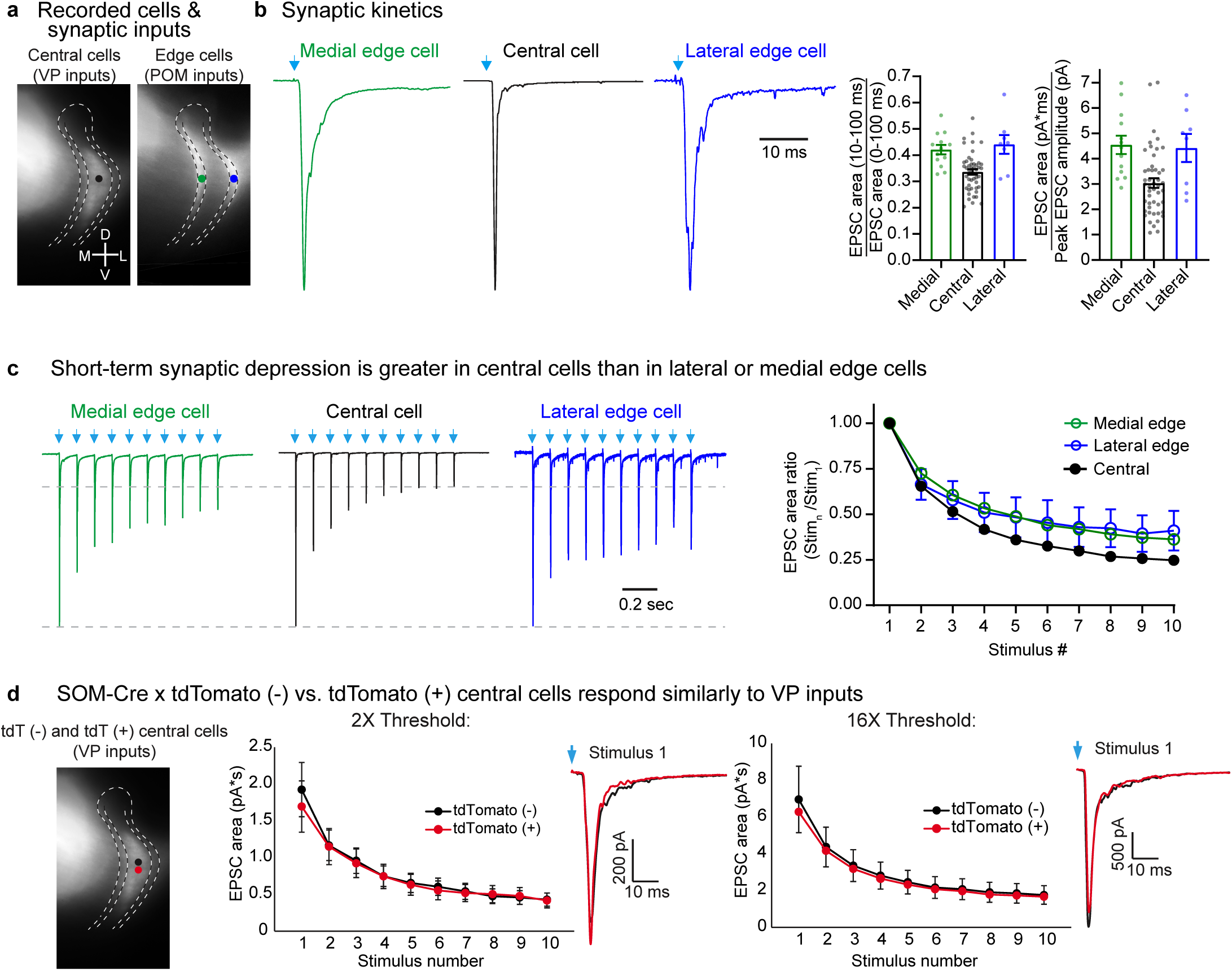
Comparison of EPSC kinetics and short-term synaptic depression based on topography or reporter labeling. **a**, Schematic showing approximate locations of TRN somata targeted for each subcircuit. EPSCs in medial and lateral edge cells were examined separately here, using the dataset from Fig. 3 a-c. V-clamp at -84 mV. **b**, Kinetics of the EPSCs. Left, optically evoked EPSCs of example cells from each group (stimulus 1 of (c)). EPSCs evoked by POM inputs to medial and lateral edge cells were broader, with greater fractional areas in the later portions of the responses than EPSCs evoked by VP inputs to central cells. Right, kinetic differences between responses are shown for two methods of quantification. The first is the ratio of the EPSC area during the last 90 ms of the response (10-100 ms from stimulus onset) to the full EPSC area (0-100 ms from stimulus onset; left graph). This ratio, which describes the fractional area of the late portion of the response, varied among cells groups (ANOVA, p<0.0001). Central values were significantly lower than medial and lateral edge values (two-tailed Bonferroni’s t-tests, p<0.002 and p<0.002, respectively), whereas the two edge groups were not different from one another (two-tailed Bonferroni’s t-test, p=0.99). As a second method of quantifying response kinetics we calculated the ratio of the full EPSC area (0-100 ms from stimulus onset) to the peak EPSC amplitude (right graph – as in Fig. 3b). This ratio also varied as a function of cell group (ANOVA, p<0.0005). Again, central values were significantly lower than medial and lateral edge values (two-tailed Bonferroni’s t-tests, p<0.002 and p<0.03, respectively), whereas the two edge groups were not different from one another (two-tailed Bonferroni’s t-test, p=0.99). The half-widths of EPSCs were not significantly different between any of the groups (all p>0.05 for two-tailed Bonferroni’s t-tests; data not shown). For panels b-c, central cells: 48 cells from 14 mice; medial cells: 13 cells from 11 mice, lateral cells: 8 cells from 7 mice. **c**, Left, representative traces illustrating the short-term synaptic depression of optically evoked EPSCs across groups (10 Hz trains of LED stimuli: arrows). Right, average EPSC areas measured during the first 10 ms of the responses. Depression of VP-evoked responses in central cells was stronger than depression of POM evoked responses in medial or lateral cells (two-tailed Bonferroni’s t-tests; central vs. lateral and central vs. medial, both p<0.0001), whereas depression in the two edge cell groups did not differ significantly (two-tailed Bonferroni’s t-test, p=0.99). **d**, Left, schematic showing experimental design. Recordings were made in slices from SOM-Cre X Ai14 mice. ChR2-eYFP was virally expressed in VP cells. We optically stimulated the VP cells/axons and recorded evoked synaptic responses from closely spaced tdTomato (tdT)-positive and tdT-negative neurons in the central zone of TRN. We ensured that both cells had similar eYFP terminal labeling surrounding them; usually the cells were separated by less than 50 µm. The goal was to test whether there were response differences between central TRN cells that were SOM-Cre x tdT-positive versus -negative. Neurons were tdT-positive if they expressed Cre recombinase at any stage of their lives; in SOM-Cre mice, TRN central zone neurons appeared to express Cre only early in development (see main text; Fig. 1b, e; Extended Data Figs. 1a, 2,3). Middle and right, average optically evoked synaptic responses (1 ms LED pulses, 10 Hz trains) for tdT-positive and tdT-negative cell groups for stimuli delivered at 2X (middle) and 16X (right) the threshold for evoking a clear EPSC. Threshold intensity was not significantly different between tdT-positive and tdT-negative cells. The average LED power used at 2X threshold was 2.13 mW. Response magnitudes, short-term EPSC depression, and kinetics (inset traces) did not differ between tdT-positive and -negative cells at 2X threshold (n=19 pairs of cells) or at 16X threshold (n=12 pairs of cells) (ANOVA, p=0.98 and p=0.61, respectively; all p>0.05 for cell-type comparisons on individual stimuli in the trains, two-tailed Bonferroni’s t-tests). Thus, VP synaptic inputs appear to be no different for central TRN cells that express SOM-Cre during early development than for those that are SOM-Cre negative continuously. Data expressed as mean ± SEM.

**Extended Data Figure 6.**
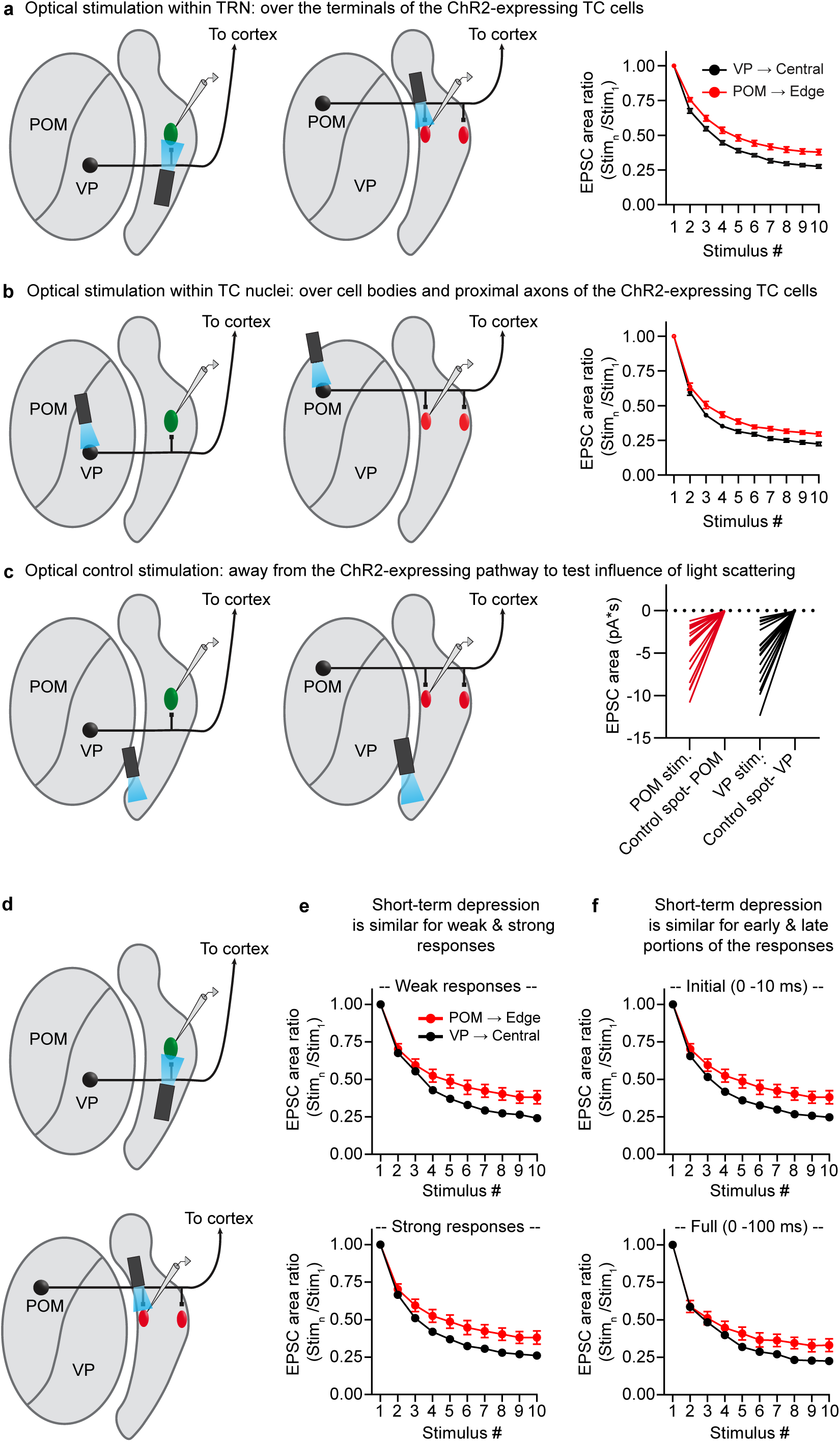
Differences in short-term depression between the two somatosensory thalamoreticular pathways do not depend on axon terminal stimulation, stimulus strength, or the temporal components of the responses measured. Some studies have shown that synaptic release probability and short-term plasticity can be abnormal when optogenetic light stimuli are directed at opsin-expressing presynaptic terminals^47^. We addressed this issue in the experiments described in panels a-c. **a**, VP synapses onto central TRN cells undergo greater short-term depression than POM synapses onto edge cells when optically stimulating the thalamocortical (TC) terminals within TRN. Left, schematic showing recording configuration when optically stimulating over the terminals of the ChR2-expressing TC cells. For the experiments of (a-c), GABA_A_ receptors and T-type calcium channels were blocked with picrotoxin (50 µM) and TTA-P2 (1 µM), respectively. Right, average short-term synaptic depression. Light intensities were chosen to evoke ∼1.5 - 3 nA peak responses. EPSC areas were measured over the first 10 ms of each response. There was greater short-term depression for the VP→Central cell pathway (n=20 cells, 5 mice) than the POM→Edge cell pathway (n=17 cells, 8 mice) (p < 0.0001, ANOVA stim. 2-10; all p < 0.005 from stimulus 5-10, two-tailed Bonferroni’s t-test). **b**, VP inputs to central cells remain more depressing than POM inputs to edge cells when stimulating within the TC nuclei. Left, schematic showing recording configuration when optically stimulating over cell bodies and proximal axons of the ChR2-expressing TC cells. Same cells as panel a. Right, average short-term synaptic depression was greater in central than edge cells (p < 0.0001, ANOVA stim. 2-10; all p < 0.05 from stimulus 5, 7-10, two-tailed Bonferroni’s t-test). This pattern of short-term depression in central cells (VP inputs) and edge cells (POM inputs) did not differ from the previously observed pattern in the same group of cells when stimulating over the TC terminals (panel a) or in other experiments of the study (d-e, Fig. 3, Extended Data Fig. 5). **c**, Optical control indicating that light scattering and spread to the TC terminals did not trigger the responses observed when stimulating within TC nuclei (i.e. the responses in panel b). Left, schematic showing recording configuration. The LED was directed to ventrally located ‘control spots’, away from the ChR2-expressing TC pathway but equally distant from the TC terminals as the effective stimulus spots within the TC nuclei. Therefore, if synaptic responses were observed, they would be triggered by light spreading to the ChR2-expressing TC terminals within the TRN. The same cells were recorded as in panels a-b. Right, EPSC areas when stimulating over the TC nuclei vs. the control spots using matching LED parameters (intensities and spot sizes). No clear responses were evoked when stimulating over the control spots, suggesting that light spread to the terminals did not drive the responses when stimulating within the TC nuclei (all p < 0.0001; two-tailed paired t-tests). **d**, Schematic showing recording configuration for panels e-f when optically stimulating VP inputs to central cells and POM inputs to edge cells. ChR2-expressing VP and POM inputs were photostimulated at their terminals (1 ms LED pulse, 10 Hz stimuli) and EPSCs were recorded in central and edge TRN cells, respectively. **e**, Differences in short-term depression of central and edge cells does not change with varying input strength. Maximum response amplitudes tended to be larger for central than for edge cells. Here we compared short-term depression of synaptic responses in central and edge cells, varying stimulus intensities for the central cells so that their responses were either weaker (top) or stronger (bottom) than those of edge cells (mean central EPSCs: 1.3 ± 0.2 nA and 5.5 ± 0.8 nA, respectively; mean edge EPSCs 2.1 ± 0.3 nA). There were no major differences in patterns of short-term synaptic depression for the weaker versus stronger responses. In both cases, there was more depression for the central cells than the edge cells (p < 0.0001, ANOVA stim. 2-10; all p < 0.05 from stimulus 5-10, two-tailed Bonferoni’s t-test; central weak responses: n=45 cells, 11 mice; central strong responses: n=32 cells, 9 mice; edge cells 21 cells, 14 mice). Central responses were acquired at ∼2X response threshold for the top row (average LED power=1.92 mW) and ∼16X response threshold for the bottom row (average LED power=12.5 mW). **f**, Short-term synaptic depression is similar whether EPSCs are measured only during the initial fast components of the responses (the first 10 ms) or across longer durations (100 ms). Top, normalized EPSC area dynamics for the initial 10 ms of the responses (same plot shown on Fig. 3c). The EPSC amplitudes were similar for central and edge cells to stimulus 1 (mean central EPSCs: 2.0 ± 0.2 nA; mean edge EPSCs 2.1 ± 0.3 nA, unpaired two-tailed t-test, p=0.93). There was greater short-term depression for the central cells than the edge cells (p < 0.0001, ANOVA stim. 2-10; all p < 0.005 from stimulus 5-10, two-tailed Bonferroni’s t-test; central cells: 48 cells, 14 mice; edge: 21 cells, 14 mice). Bottom, normalized area dynamics for the full responses (0-100 ms). Again, there was greater short-term depression for the central cells than the edge cells (p < 0.0001, ANOVA stim. 2-10; p < 0.05 for stimulus 8-10, two-tailed Bonferroni’s t-test). Thus, the overall short-term synaptic depression patterns are similar for early and later components of the EPSCs. Data expressed as mean ± SEM.

**Extended Data Figure 7.**
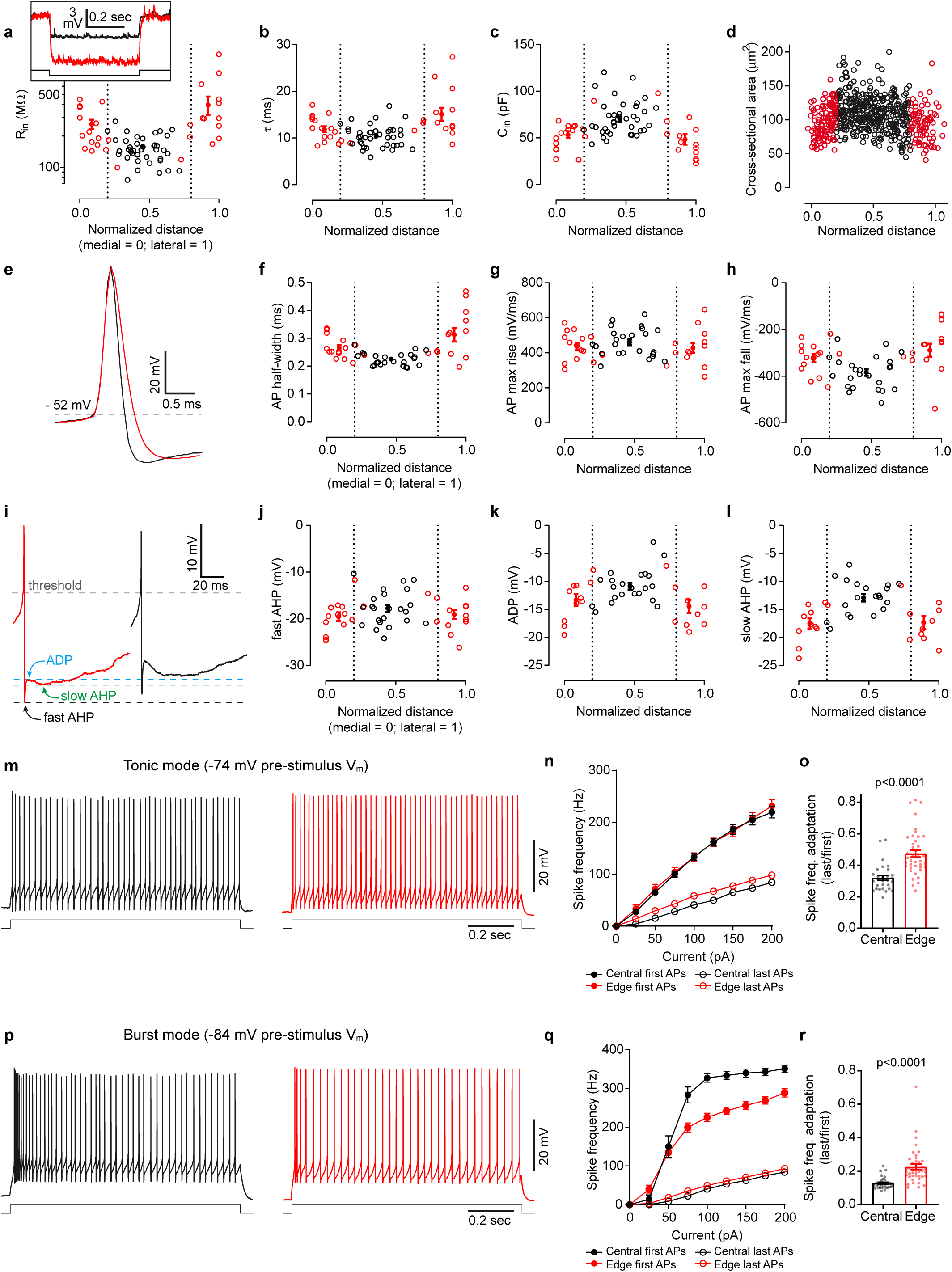
Intrinsic cellular properties vary according to medial-lateral position in the TRN. **a-c**, Inset, voltage responses to hyperpolarizing current steps for representative central and edge cells. Throughout this figure, central cells are colored black and edge cells are red. Note the higher input resistance for the edge cell (316 vs. 152 MΩ). Plots show passive membrane properties for central and edge cells as a function of soma position (far medial edge = position 0, far lateral edge = position 1; steady-state V_m_ ∼ -84 mV). Open circles indicate individual cell measurements. Filled circles are averages for the cell groups: medial edge, central, and lateral edge. **d**, Central cells have larger somas than edge cells. ROIs outlining complete somas were hand-drawn with the aid of ROIManager (ImageJ) using the PV-IHC tissue from Fig. 1 (n=575 cells; 3 sections; 3 mice). The mean soma area for central and edge were 110.05 µm^2^ vs 95.59 µm^2^, respectively (unpaired two-tailed t-test, p<0.0001). Cell areas appear to change gradually with location. **e**, Representative examples of edge cell and central cell action potentials (APs). **f-h**, AP properties as a function of soma location. Half-width (f), AP maximum rate-of-rise (g), AP maximum rate-of-repolarization (h). Edge cells have wider APs due to slower repolarization slopes, suggesting differences in potassium conductances. **i**, Representative AP afterpotentials of each cell type. Edge cells generally have more hyperpolarized fast-afterhyperpolarizations (fast-AHPs), afterdepolarizations (ADPs), and slow-afterhyperpolarizations (slow-AHPs). **j-l**, fast-AHP (j), ADP (k), and slow-AHP (l) as a function of somatic location (APs evoked from a steady state V_m_ ∼ -74 mV; values relative to AP threshold). **m**, Representative examples of repetitive tonic firing evoked by 75 pA current steps (from -74mV baseline potentials) for central and edge cells. **n**, Spike Frequency vs. Current Intensity (FI) plots. Closed circles show the frequency for the first 2 spikes in each train. Open circles show the frequency for the last 2 spikes. The initial firing frequency is similar between edge and central cells at multiple current steps. However, edge cells fire at higher frequencies than central cells by the end of the current pulses, perhaps because edge cells have higher input resistances (for panels n-o, central cells n= 29, edge cells n=40). **o**, Central cells have more spike frequency adaptation than edge cells (unpaired two-tailed t-test p<0.0001). **p-r**, Same as panels m-o, except responses were evoked from steady-state potentials of -84 mV to elicit intrinsic bursts. Central cells showed more pronounced bursts than edge cells. On average, central cells required stronger currents to evoke spiking than edge cells when stimulated from -84 mV steady-state potentials (q) (see Supplementary Information 1 for more threshold information). However, the initial spike frequencies for central cells were much higher than those for edge cells due to their bursty nature (q). The frequencies by the ends of the trains were nearly identical for central and edge cells (q). Central cells had stronger spike frequency adaptation than edge cells (r), largely due to stronger initial bursting in central cells (unpaired two-tailed t-test, p<0.0001; for panels q-r: central cells n= 31, edge cells n=42). For the physiological measurements in this figure, central and edge cell recordings were interleaved within slices (drawn from at least 8 mice per measured property). Data expressed as mean ± SEM. See Supplementary Information 1 for additional intrinsic physiological properties and statistical comparisons.

**Extended Data Figure 8.**
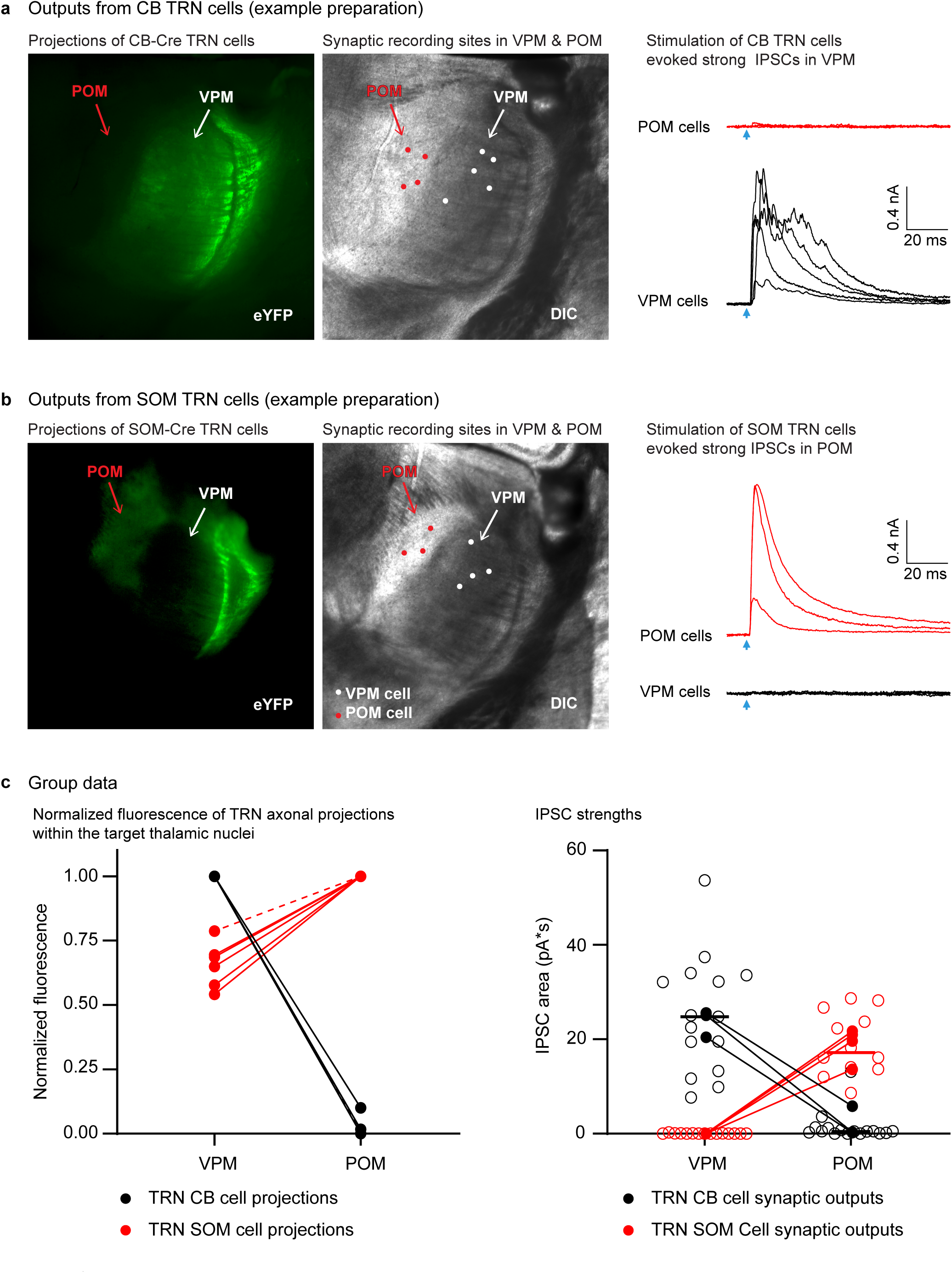
Inhibitory output from the TRN: SOM-expressing TRN edge cells project to POM, whereas CB cells project to VPM. **a**, Fluorescence (left) and DIC (middle) images from a live 300 µm slice following ChR2-eYFP expression in CB-Cre-positive TRN cells. Fluorescent axons of these TRN CB cells target VPM (Ventral Posterior Medial nucleus) but not POM. White and red circles show the locations of recorded VPM and POM neurons, respectively. Right, strong IPSCs were evoked in the VPM neurons (black) whereas virtually no IPSCs were evoked in the POM neurons (red) by optical activation of CB-Cre TRN cells (each trace is the response for one cell, averaged over 10 sweeps; V_hold_ -34 mV). Arrows indicate onsets of LED flashes (1 ms duration). Glutamate receptors were blocked by applying APV (50 µM) and DNQX (20 µM). The experiment was replicated in 4 slices from 3 CB-Cre mice. **b**, Same as (a) except ChR2-eYFP expression was in SOM-Cre TRN edge cells. Notice both the strong axonal targeting and IPSCs in POM, rather than in VPM. The experiment was replicated in 6 slices from 5 SOM-Cre mice. **c**, Left, group plot comparing fluorescence projection intensities to VPM and POM from the two TRN cell types. Fluorescence was normalized to the maximum level across the thalamic relay nuclei, with zero values set to the minimum level in the slice (n=6 slices from 5 SOM-Cre mice; n=4 slices from 3 CB-Cre mice; dashed lines represent a second slice for one mouse). Projections of CB-Cre TRN cells were stronger in VPM than in POM, whereas projections of SOM-Cre TRN cells were stronger in POM than in VPM (all P<0.0001, two-tailed Bonferroni’s t-tests). Right, strengths of IPSCs recorded in VPM and POM reflected their anatomical inputs from TRN. IPSCs evoked by photostimulation of CB-Cre TRN central cells (black) were large in VPM (n=15, 3 mice) and very weak in POM (n=16, 3 mice) (two-tailed Bonferroni’s t-test, p<0.0001). Conversely, IPSCs evoked by stimulation of SOM-Cre edge cells (red) were strong in POM (n=12 cells, 4 mice) and undetectable in VPM (n=15 cells, 4 mice) (two-tailed Bonferroni’s t-test, p<0.0001). Open circles show response areas for individual cells. Closed circles show averages per mouse. Comparisons were made using matched LED power. Data expressed as mean ± SEM.

**Extended Data Figure 9.**
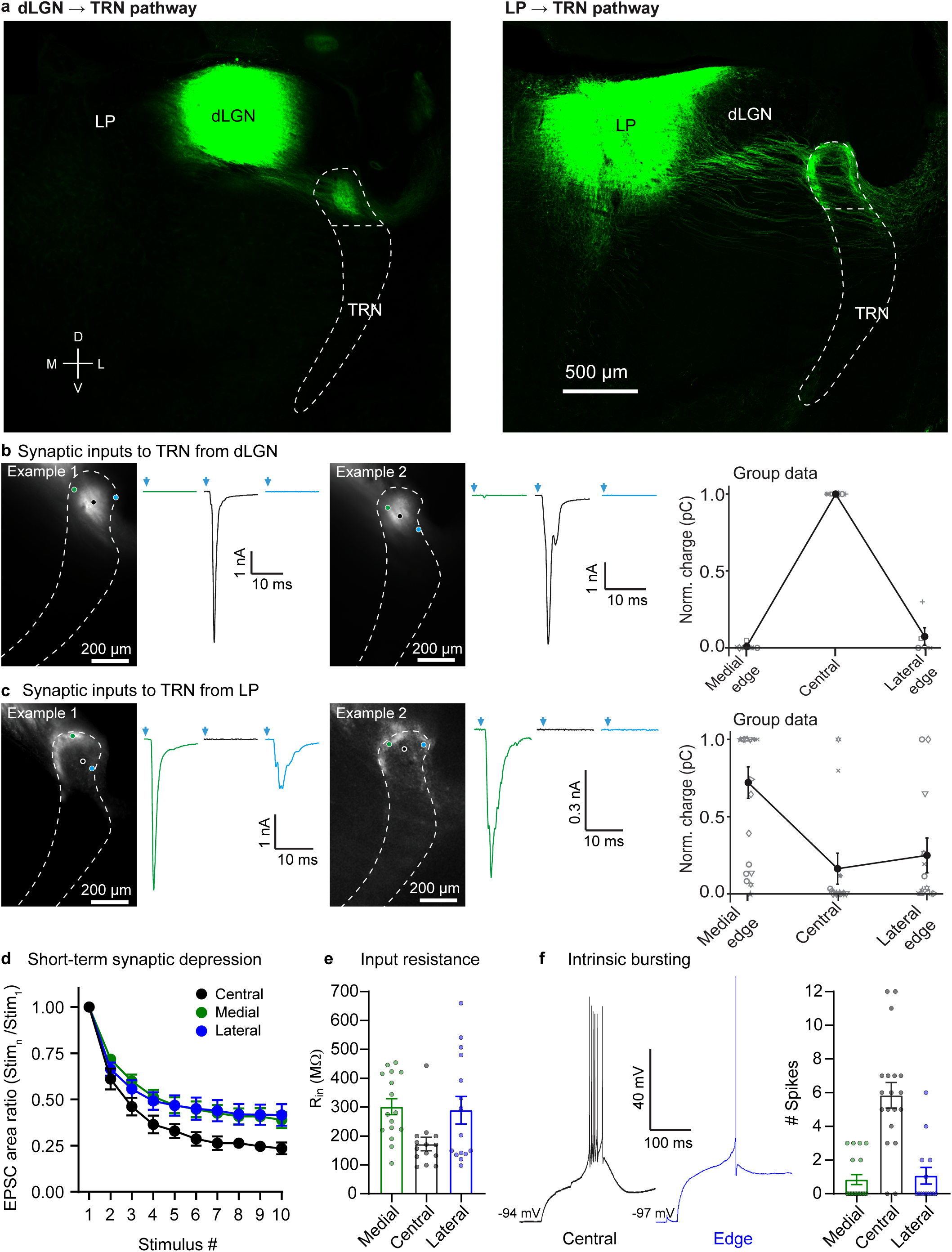
Visual TRN and somatosensory TRN have similar center and edge subcircuits. **a**, Representative images showing dLGN and LP injection sites and anterograde projections to TRN. dLGN projects to the central zone whereas LP projects to the edges of the visual sector of TRN. This experiment was replicated in 5 mice with dLGN viral injections, and in 13 mice with LP viral injections. **b**, Two representative examples (left and middle panels) of synaptic responses from cells across the medial-lateral axis of visual TRN evoked by photostimulation of dLGN. Cells in the center of visual TRN received strong input from dLGN, whereas inputs to cells located near the edges of TRN were very weak or undetectable (average of 10 sweeps, -84 mV). Right, relationship between soma location in TRN and synaptic response amplitude (n=19 cells, 6 slice preparations, each preparation represented by a different symbol shape; means for the 3 zones are represented by filled symbols and connected by lines), showing that cells in the center of visual TRN respond strongly to dLGN input and those in the medial and lateral edges have minimal dLGN inputs (as in Fig. 2, and Extended Data Fig. 4). **c**, Same as (b) except characterizing synaptic inputs to TRN from LP. Cells along the medial edge (and in some cases along the lateral edge) of visual TRN received strong input from LP. In contrast, cells located in the center of visual TRN generally had weak or no responses to LP input (average of 10 sweeps, -84 mV; n=49 cells,12 preps). **d**, Dynamic properties of visual thalamoreticular responses. To study short-term synaptic depression, photostimulation intensities were chosen to evoke ∼2.5 nA peak responses (average LED powers: 20.8 mW for LP, 6.6 mW for dLGN). EPSC areas were measured over the first 10 ms of each response. Short-term depression was more pronounced for the central cells than the edge cells (p < 0.0001, ANOVA, stim. 2-10). Depression in the two edge cell groups did not differ (two-tailed Bonferroni’s t-test, all p>0.99). Depression of dLGN-evoked responses in central cells was stronger than depression of LP-evoked responses in medial or lateral cells (two-tailed Bonferroni’s t-test; central vs. medial, central vs. lateral, both p<0.0001; central group: 7 cells, 5 mice; medial group: 17 cells, 11 mice; lateral group: 7 cells, 7 mice). Analogous to subcircuits of the somatosensory TRN, dLGN-evoked EPSCs of central cells in the visual TRN were less prolonged than LP-evoked EPSCs of edge cells (not shown). For example, the ratio of the EPSC areas to the peak EPSC amplitudes (as in Fig. 3b) varied as a function of cell group (ANOVA, p<0.009; central cells ratio = 2.74 ± 0.24; medial cells ratio = 5.27 ± 0.49; lateral cells ratio = 5.14 ± 0.60, same cells as for short-term depression analysis, above). The two edge groups did not differ (two-tailed Bonferroni’s t-test, p=0.99). Central cell ratios were significantly lower than medial and lateral cell ratios (two-tailed Bonferroni’s t-test, p<0.009, <0.05, respectively). **e**, Input resistances of medial edge (n=16 cells, 14 mice), central (n=14 cells, 13 mice), and lateral edge cells (n=15 cells, 13 mice) varied as a function of cell group (p<0.03, ANOVA). The two edge groups were not different from one another (two-tailed Bonferoni’s t-test, p=0.99). Central cell input resistance was significantly different from medial, although not from lateral (two-tailed Bonferroni’s t-tests, p<0.04 and p=0.074, respectively). **f**, Central cells of visual TRN had more robust intrinsic bursting than those along the edges. Left, representative stimulus-offset bursts. Right, average number of spikes during bursts was 5.8 ± 0.76 for central cells (n=19 cells, 15 mice), 0.8 ± 0.30 for medial cells (n=19 cells, 14 mice), and 1.1 ± 0.50 for lateral cells (n=14 cells, 12 mice) (p<0.0001, ANOVA). The two edge groups were not different from one another (two-tailed Bonferoni’s t-test, p=0.99), but the edge cells discharged far fewer spikes per burst than the central cells (two-tailed Bonferoni’s t-test, p<0.0001 for both medial vs. central cells and lateral vs. central cells). For the experiments represented in this figure, the boundaries of the TRN were determined using either tdTomato expression driven by SOM-Cre or PV-Cre, IHC for PV, or brightfield images. Data expressed as mean ± SEM.

**Extended Data Figure 10.**
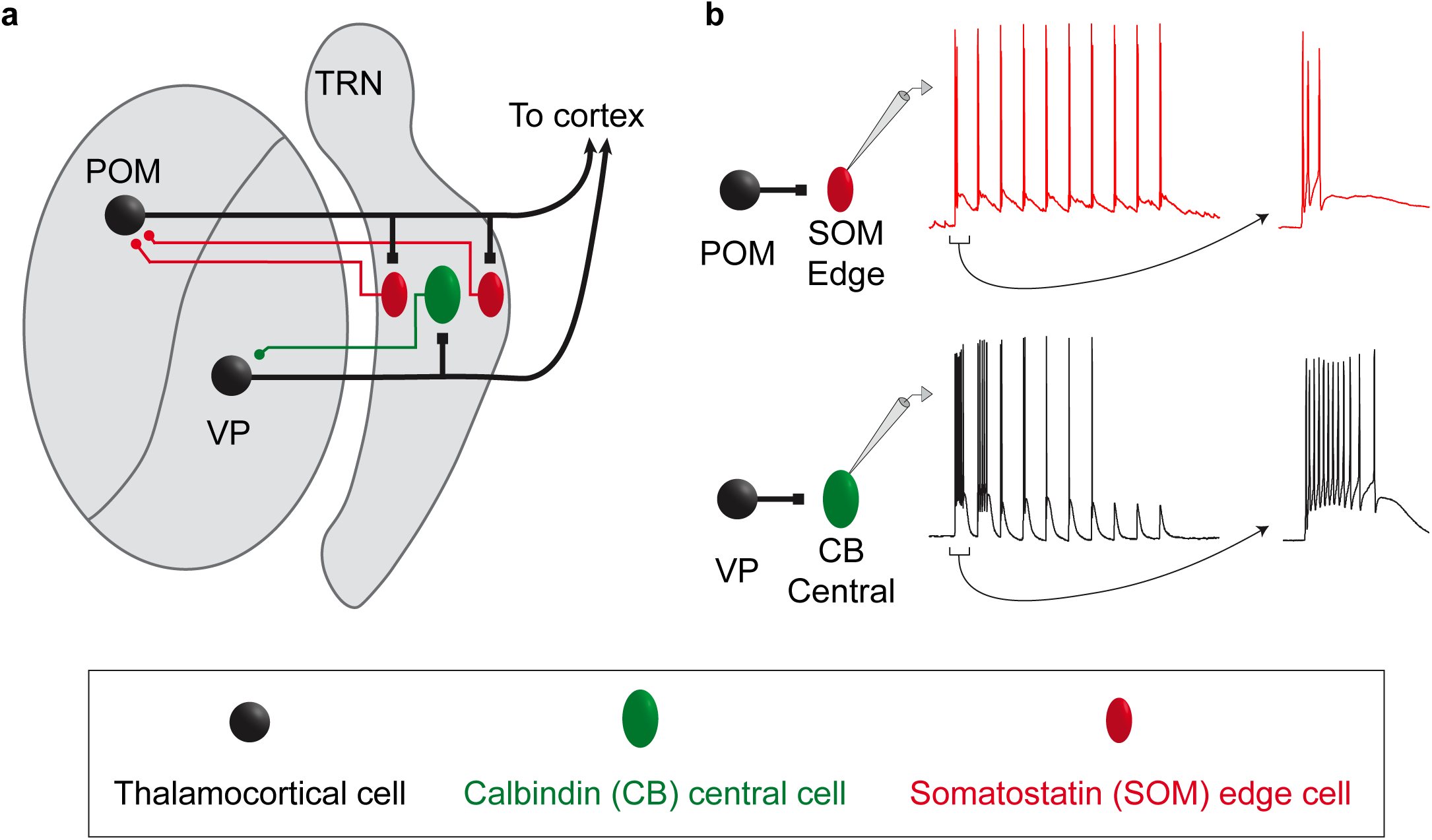
Organizational and operational scheme for somatosensory circuits of the TRN. **a**, Schematic illustrating the organization of the somatosensory sector of TRN. Excitatory inputs from the primary VP thalamic nucleus innervate the CB-dense central zone of TRN (green cells), whereas higher-order POM inputs innervate the SOM-dense edges (red cells). The inhibitory projections of TRN cells to VP and POM appear to reciprocate the patterns of their inputs (colored axons in the schematic). The spatial organization of the neurochemically distinct TRN cell types, and the stark segregation of their primary and higher order connections, suggest a core-shell organization for somatosensory circuits of the TRN. **b**, Distinct responses of central and edge TRN cells to their thalamic inputs. The spiking responses of central and edge cells are determined by the dynamic synaptic properties of their respective thalamic inputs combined with the intrinsic physiological properties of the TRN cells themselves. The spiking of central cells evoked by VP input is phasic, with strong initial bursts that deeply depress with repetitive stimulation; this is consistent with a core circuit that processes transient sensory signals. In contrast, the spiking responses of edge cells to POM input are more sustained, with initially weaker but more persistent activation; this shell-like circuit is suited to processing temporally distributed higher-order signals. TRN: thalamic reticular nucleus; POM: posterior medial nucleus; VP: ventral posterior nucleus

Supplementary Information 1

Intrinsic physiological properties of central and edge TRN neurons

| Somatosensory TRN |  |  |  |  |  |  |
| --- | --- | --- | --- | --- | --- | --- |
| | Burst Mode (-84 mV pre-stimulus $V_m$) | | | Tonic Mode (-74 mV pre-stimulus $V_m$) | | |
|  | Central Cells | Edge Cells | P values | Central Cells | Edge Cells | P values |
| Resting potential (mV) | $-83 \pm 0.3$<br>(157) | $-85 \pm 0.4$<br>(101) | <0.0001 | n/a | n/a | n/a |
| Input resistance (MΩ) | $157 \pm 7.1$<br>(42) | $316 \pm 40.5$<br>(31) | <0.0001 | $278 \pm 24.3$<br>(33) | $495 \pm 59.5$<br>(32) | <0.001 |
| Membrane time constant (ms) | $10.4 \pm 0.34$<br>(42) | $13.3 \pm 0.76$<br>(31) | <0.0005 | $17.5 \pm 1.16$<br>(33) | $20.5 \pm 1.22$<br>(32) | 0.08 |
| Input capacitance (pF) | $70 \pm 2.8$<br>(42) | $51 \pm 3.2$<br>(31) | <0.0001 | $67 \pm 2.6$<br>(33) | $49 \pm 2.8$<br>(32) | <0.0001 |
| Rheobase (pA) | $61 \pm 4.5$<br>(25) | $39 \pm 4.1$<br>(28) | <0.0005 | $24 \pm 3.3$<br>(26) | $16 \pm 2.4$<br>(28) | <0.05 |
| AP threshold (mV) | $-58 \pm 0.7$<br>(25) | $-56 \pm 0.8$<br>(28) | <0.05 | $-56 \pm 0.7$<br>(26) | $-55 \pm 0.7$<br>(28) | 0.4 |
| AP amplitude (mV) | $69 \pm 1.6$<br>(22) | $72 \pm 1.7$<br>(23) | 0.2 | $68 \pm 1.5$<br>(24) | $69 \pm 1.8$<br>(28) | 0.4 |
| AP maximum rate of rise (mV/ms) | $488 \pm 17.8$<br>(22) | $481 \pm 18.6$<br>(23) | 0.8 | $459 \pm 16.4$<br>(24) | $434 \pm 17.1$<br>(28) | 0.3 |
| AP maximum rate of fall (mV/ms) | $-400 \pm 17.4$<br>(22) | $-336 \pm 15.4$<br>(23) | <0.01 | $-384 \pm 14.1$<br>(24) | $-307 \pm 15.8$<br>(28) | <0.0005 |
| AP half-width (ms) | $0.22 \pm 0.005$<br>(22) | $0.27 \pm 0.010$<br>(23) | <0.0005 | $0.23 \pm 0.005$<br>(24) | $0.29 \pm 0.013$<br>(28) | <0.0001 |
| Fast afterhyperpolarization (fast AHP) (mV) | $-14.5 \pm 0.71$<br>(22) | $-16.1 \pm 0.82$<br>(22) | 0.2 | $-17.7 \pm 0.78$<br>(23) | $-19.5 \pm 0.72$<br>(23) | 0.09 |
| Afterdepolarization (ADP) (mV) | *na* | *na* | *na* | $-10.8 \pm 0.64$<br>(22) | $-13.9 \pm 0.80$<br>(21) | <0.005 |
| Slow afterhyperpolarization (slow AHP) (mV) | $-18.2 \pm 0.64$<br>(22) | $-20.1 \pm 0.82$<br>(22) | 0.08 | $-12.9 \pm 0.67$<br>(20) | $-17.4 \pm 0.72$<br>(21) | <0.0001 |
| Offset burst # APs | $9.6 \pm 0.85$<br>(34) | $2.3 \pm 0.51$<br>(38) | <0.0001 | *na* | *na* | *na* |
| Offset burst peak frequency (Hz) | $364 \pm 23.8$<br>(34) | $108 \pm 23.2$<br>(38) | <0.0001 | *na* | *na* | *na* |
| Offset burst mean frequency (Hz) | $243 \pm 15.4$<br>(34) | $87 \pm 18.6$<br>(38) | <0.0001 | *na* | *na* | *na* |
| Spike frequency adaptation (Last/First) | $0.13 \pm 0.01$<br>(31) | $0.23 \pm 0.02$<br>(42) | <0.0001 | $0.32 \pm 0.02$<br>(29) | $0.47 \pm 0.02$<br>(40) | <0.0001 |

Intrinsic physiological properties of central and edge TRN neurons

| Visual TRN |  |  |  |  |  |  |
| --- | --- | --- | --- | --- | --- | --- |
| | Burst Mode (-84 mV pre-stimulus $V_m$) | | | Tonic Mode (-74 mV pre-stimulus $V_m$) | | |
|  | Central Cells | Edge Cells | P value | Central Cells | Edge Cells | P value |
| Resting potential (mV) | $-84 \pm 0.3$<br>(21) | $-85 \pm 0.4$<br>(42) | 0.47 | *na* | *na* | *na* |
| Input resistance (MΩ) | $173 \pm 23.3$<br>(14) | $296 \pm 26.8$<br>(31) | <0.01 | *na* | *na* | *na* |
| Membrane time constant (ms) | $10.9 \pm 0.75$<br>(14) | $12.4 \pm 0.53$<br>(31) | 0.12 | *na* | *na* | *na* |
| Input capacitance (pF) | $70 \pm 4.3$<br>(14) | $51 \pm 5.7$<br>(31) | <0.05 | *na* | *na* | *na* |
| Offset burst # APs | $5.8 \pm 0.76$<br>(19) | $0.9 \pm 0.27$<br>(33) | <0.0001 | *na* | *na* | *na* |
| Offset burst peak frequency (Hz) | $207 \pm 23.1$<br>(19) | $32 \pm 11.3$<br>(33) | <0.0001 | *na* | *na* | *na* |
| Offset burst mean frequency (Hz) | $107 \pm 8.0$<br>(19) | $25 \pm 13.7$<br>(33) | <0.0001 | *na* | *na* | *na* |

**Supplementary Information 1. Intrinsic physiological properties of central and edge TRN cells**.

Passive membrane properties, action potential (AP) kinetics, and repetitive firing properties are shown for central and edge neurons of somatosensory TRN (page 1) and visual TRN (page 2). Properties were measured in current-clamp mode, with steady-state Vm set to either -84 mV (“burst mode”, left) or -74 mV (“tonic mode”, right). “Offset bursts” are intrinsic bursts triggered by the offset of hyperpolarizing current pulses, as illustrated in Fig. 3d and Extended Data Fig. 9f. The numbers of cells are shown in parenthesis. T-tests were unpaired and two-tailed. Data are expressed as mean ± SEM.

## Methods

### Animals

All procedures were approved by, and complied with all ethical regulations of, the Brown University Institutional Animal Care and Use Committee. The following mouse lines were used: SOM-IRES-Cre (The Jackson Laboratory, 013044), PV-Cre (The Jackson Laboratory, 008069), Calb1-IRES-Cre-D (“CB-Cre”; The Jackson Laboratory, 028532), Vgut2-IRES-Cre (The Jackson Laboratory, 016963), GPR26-Cre (STOCK Tg(Gpr26-cre)KO250Gsat/Mmucd, MMRRC), Ai14 (The Jackson Laboratory, 007908), ICR (Charles River, CD-1[ICR], Strain Code 022). To fluorescently target somatostatin (SOM) or parvalbumin (PV) cells in TRN, we bred the respective homozygous Cre mice (SOM-IRES-Cre, PV-Cre) with homozygous Cre-dependent tdTomato reporter mice (Ai14). In some experiments we instead injected viruses carrying Cre-dependent GFP genes into Cre mice (specified in the descriptions of the individual experiments, below). Of the 124 mice used in this study, 91 had C57 genetic backgrounds, 10 had ICR genetic backgrounds, and 23 had mixed C57/ICR backgrounds. Mice were maintained on a 12:12-hr light-dark cycle, group-housed, and provided food and water ad libitum.

### Immunohistochemistry

Mice from postnatal ages 22-26 days were deeply anesthetized with Beuthanasia-D and intracardially perfused with 0.1 M phosphate buffer (PB) followed by 4% paraformaldehyde (in PB). Brains were post-fixed overnight at 4°C in the same fixative and then transferred to a 30% sucrose/0.1 M PB solution until sectioning (4°C, 2-3 days). Forty µm-thick brain sections were cut on a freezing microtome at a somatosensory thalamocortical plane (35° tilt from coronal) ^46^ designed to contain the somatosensory thalamus, TRN, barrel cortex and many of their interconnections. Next the sections were immunostained. Briefly, they were washed 5 times in 0.1 M phosphate buffer containing 0.15 M NaCl, pH 7.4 (PBS) (5 min per wash), pre-incubated for 2 h at room temperature with a blocking solution (10% normal goat serum, 2% Triton X-100, 0.1% Tween 20 in 0.1 M PB), then incubated with primary antibodies for 5 days at 4°C. After the primary incubation, sections were washed 8 times in PBS (5 min per wash), pre-incubated for 2 h in blocking solution, incubated with a secondary antibody solution for 3 days at 4°C, then washed 8 times in PBS and 3 times in PB (5 min per wash). Sections were mounted and cover-slipped using Prolong Gold or Prolong Gold with DAPI (Molecular Probes P36930 or P36931). Primary antibodies were: mouse monoclonal anti-parvalbumin (1:1000, Swant clone 235, Lot 10-11F), rabbit polyclonal anti-Calbindin D-28k (1:1000 Swant clone CB-38a, Lot 9.03), mouse monoclonal anti-Calbindin D-28k (1:1000 Swant clone 300, Lot 07(F)), mouse monoclonal anti-NeuN (1:1000, Millipore MAB377, Clone A60, Lot 2549411). Secondary antibodies were goat anti-mouse IgG (H+L) cross-adsorbed secondary antibody Pacific Blue (1:250, Molecular Probes P31582), goat anti-rabbit IgG (H+L) cross-adsorbed secondary antibody Alexa Fluor 647 (1:500, Molecular Probes A21244), goat anti-mouse IgG (H+L) cross-adsorbed secondary antibody Alexa Fluor 488 (1:250, Molecular Probes A11001), goat anti-rabbit IgG (H+L) highly cross-adsorbed secondary antibody Alexa Fluor 488 (1:250, Molecular Probes A11034). To test the specificity of the antibodies, no-primary and no-secondary controls were conducted for each. In addition, for the calbindin antibodies, we performed assays in which the antibodies were pre-adsorbed with calbindin protein (3.33 µg/1 µl of antibody, Swant recombinant rat Calbindin D-28) before tissue incubation. There was no clear labeling under any of the control conditions, providing support for the effectiveness and specificity of the antibodies. Confocal image stacks were taken on a Zeiss LSM 800, 20X objective, 0.26 µm pixel diameter.

For analysis of the immunohistochemical material, we selected sections spanning the somatosensory sector of TRN (approximately -1.7 mm to -1.1 mm from bregma), generally focusing on the anterior-posterior center of that range (i.e., -1.4 mm from bregma). The CellCounter ImageJ plugin was used to count cells positive for the tested markers. For accurate alignment across subjects and sections, small differences in TRN shapes were minimized by warping them to an average reference TRN image using the bUnwarpJ plugin in ImageJ. At least 12 landmark points were applied around the outer boundaries of each TRN (evenly spaced, 100 µm between points) for alignment. The TRN boundaries were identified using either the SOM-Cre x tdTomato or PV-Cre x tdTomato channel. Resulting corrections were applied to all channels.

### Fluorescence *in situ* hybridization (FISH)

Mice (P21-P27) were deeply anesthetized with isofluorane and decapitated. The brains were removed while submerged in 4°C saline solution and fresh-frozen on liquid nitrogen. Frozen brains were sectioned (18 µm thick) at the somatosensory thalamocortical plane (described above) using a cryostat (Leica), adhered to SuperFrost Plus slides (VWR, 48311-703), and refrozen (−80°C) until used. Samples were fixed (4% PFA), processed as instructed in the ACD RNAScope Multiplex Fluorescent v2 Assay, and then immediately incubated with DAPI and cover-slipped with ProLong Gold (Molecular Probes P36930). Probes for PV (Mm-Pvalb-C3, Cat# 421931-C3), SOM (Mm-Sst, Cat# 404631), CB (Mm-Calb1-C2 Cat# 428431-C2), and tdTomato (tdTomato-C3 cat# 317041-C3) were purchased from Advanced Cell Diagnostics. Probes were visualized using the TSA^®^ Plus Fluorescein (1:1000, Perkin Elmer NEL741E001KT), TSA^®^ Plus Cyanine 3 (1:1000, Perkin Elmer NEL744E001KT), and TSA^®^ Plus Cyanine 5 (1:1000, Perkin Elmer NEL745E001KT) evaluation kits. Confocal image stacks were taken on a Zeiss LSM 800, 20X objective, 0.26 µm pixel diameter.

One section was selected from each of 6 mice for the FISH analysis. These sections were centered on the somatosensory sector of the TRN (approximately 1.4 mm posterior to bregma). Four of the 6 mice were C57 wild-types. The remaining 2 mice were PV-Cre x Ai14 mice that expressed tdTomato in PV-Cre cells (the tdTomato-C3 probe, cat# 317041-C3, was used as a proxy for PV for these 2 mice). Images were quantified using ImageJ. Regions of interest (ROIs) were manually drawn for each cell based on the fluorescence signal of all three channels (SOM, CB, and either PV or tdTomato). For quantification, the mean fluorescence intensity for a cell’s ROI was expressed as a percentage of the intensity range across the TRN of the section: 100 x [(mean intensity for the cell’s ROI) / (Maximum Pixel Intensity in the TRN - Minimum Pixel Intensity in the TRN)]. Cells were considered positive for an RNA marker if this normalized expression for the marker was greater than 7.5%, which best matched qualitative visual assessment of expression thresholds.

Probes for PV (Mm-Pvalb-C4, Cat# 421931-C4), Cre (CRE-C3, Cat# 312281-C3), and SOM (Mm-Sst, Cat# 404631) were used to assess correspondence between Cre and SOM mRNA expression in the SOM-Cre mouse line (4 SOM-Cre x ICR Mice).

### Cell Count Analysis

For both immunohistochemical and FISH analysis, cells were counted in a region centered on somatosensory TRN extending 300 µm along the dorsal-ventral axis of the nucleus (dorsal-ventral boundaries indicated by the brackets in Fig. 1a-b, or the boxes in Extended Data Figs. 1b, 2a). This region consistently received axonal projections from S1 cortex (data not shown) and somatosensory thalamus (VP and POM; Fig. 2, Extended Data Fig. 4). Central/edge boundaries drawn at 20% and 80% of the medial-lateral distance across TRN.

### Stereotactic Injection Procedure

Mice were anaesthetized with a Ketaset-Dexdormitor mixture diluted in sterile saline (Ketaset, 70 mg/kg; Dexdormitor, 0.25 mg/kg; intraperitoneally). Once deeply anesthetized, mice were placed into a stereotactic frame, and a craniotomy was made over VP, POM, dLGN, LP, or TRN. Virus solution was then pressure-ejected into the brain via a glass micropipette attached to a Picospritzer pressure system at a maximum rate of ∼0.05 µl/min (10-30 min total injection times). Following injection, the pipette was left in place for ∼10 min before being slowly withdrawn from the brain. After surgery, mice were given Antisedan (2.5 mg/kg) to reverse the effects of Dexdormitor, and they were allowed to recover on a heating pad for ∼1 h before being returned to their home cage. Experiments were usually performed ∼ 10 days after the virus injections to allow for sufficient EGFP or opsin expression (mean: 10.4 ± 0.3, range: 8-19 days).

Adeno-associated viruses (AAVs) and lentiviruses were acquired from the University of North Carolina, Addgene or the University of Pennsylvania Vector Cores and used at the following titers: (1) rAAV2/hSyn-ChR2(H134R)-eYFP-WPREpA (titer = ∼3.93 x 10^12^ [vg]/ml), (2) pLenti-Synapsin-hChR2(H134R)-EYFP-WPRE (titer = ∼2.53 x 10^10^ [vg]/ml), (3) AAV2.Syn.DIO.hChR2(H134R)-EYFP.WP.hGH (titer = ∼2.32 x 10^13^ [vg]/ml), (4) rAAV2/hSyn-ChR2(H134R)-mCherry-WPREpA (titer = ∼2.2 x 10^12^ [vg]/ml), (5) AAV9.Syn.DIO.EGFP.WPRE.hGH (titer = ∼6.25 x 10^12^ [vg]/ml), (6) rAAV2/Syn-Flex-ChrimsonR-TdT (titer = ∼3.8 x 10^12^ [vg]/ml).

For experiments testing projections from specific thalamocortical nuclei to TRN (or from subtypes of TRN cells to thalamocortical nuclei) relatively small volumes (0.16 – 0.33 µl) of either AAV2 or lentivirus (described above) were stereotactically injected into individual presynaptic nuclei using the following mouse strains and coordinates. To test VP projections to TRN (Figs. 2-3, Extended Data Figs. 4-6), virus was injected into SOM-Cre x Ai14 (n=15), ICR (n=4), Vglut2-Cre x ICR (n=5), or PV-Cre x Ai14 (n=1) mice between postnatal days 12 and 16 (P12-16; mean = P13.1 ± 0.2 d). Average coordinates from bregma for VP were 1.99 mm lateral, -0.74 mm posterior, 3.08 mm depth. For POM projections to TRN (Figs. 2-3, Extended Data Fig. 4-6), virus was injected into SOM-Cre x Ai14 (n=8), ICR (n=6), PV-Cre x Ai14 (n=2), or GPR26-Cre x ICR (n=2) mice between P11-P14 (mean = P12.0 ± 0.3). Coordinates for POM were 1.36 mm lateral, -1.17 mm posterior, 2.86 depth. For dLGN projections to TRN (Extended Data Fig. 9), virus was injected into SOM-Cre x Ai14 (n=2), PV-Cre x Ai14 (n=2) or GPR26-Cre x ICR (n=1) mice between P14-19 (mean = P16.0 ± 0.8). Coordinates for dLGN were 2.38 mm lateral, -1.45 mm posterior, 2.35 depth). For LP projections to TRN (Extended Data Fig. 9), virus was injected into SOM-Cre x Ai14 (n=6), PV-Cre x Ai14 (n=6) or CB-Cre x Ai14 (n=1) mice between P14-17 (mean = P15 ± 0.3). Coordinates for LP were 1.63 mm lateral, -1.36 mm posterior, 2.3 mm depth. For tests of the inhibitory outputs of TRN cell subtypes to VP and POM neurons (Extended Data Fig. 8), virus was injected into SOM-Cre x ICR (n=5), CB-Cre x Ai14 (n=1) or CB-Cre x ICR (n=2) mice between P12-15 (mean = 13.7 ± 0.3). Coordinates for these TRN injections were 2.2 mm lateral; -0.53 mm posterior, 3.0 mm depth.

For assessments of the positions of TRN cell subtypes (Fig. 1e, Extended Data Fig. 1, 3), relatively large volumes (1.3-2 µl) of AAV9 (described above) were injected across the entire dorsal-ventral extent of the TRN in SOM-Cre x Ai14 (n=10), SOM-Cre x ICR (n=2) or PV-Cre x Ai14 (n=4), or ICR (n=2) mice between P14 and P25 (mean 16.3 ± 0.9). Average coordinates for these TRN injections were 2.2 mm lateral; -0.53 mm posterior, with continuous outflow of virus solution from 2.2 to 4.4 mm depth.

### Slice Preparation

Brain slices were prepared from P22-34 mice of either sex as previously described^40^. Mice were deeply anesthetized with isofluorane and decapitated. The brains were removed while submerged in cold (4°C) oxygenated slicing solution containing (in mM): 3.0 KCl, 1.25 NaH_2_PO_4_, 10.0 MgSO_4_, 0.5 CaCl_2_, 26.0 NaHCO_3_, 10.0 glucose, and 234.0 sucrose. Brains were then mounted, using a cyanoacrylate adhesive, onto the stage of a vibrating tissue slicer (Leica VT1000 or VT1200S) and somatosensory thalamocortical brain slices (300 μm thick, 35° tilt from coronal^46^) containing VP, POM, TRN, S1, and portions of dLGN and LP were obtained. Slices were incubated for ∼1 min in the cold sucrose-based slicing solution, then transferred for 20 min to a holding chamber containing warm (32°C) oxygenated (5% CO_2_, 95% O_2_) artificial cerebrospinal fluid (ACSF) solution. Finally, the slices were allowed to equilibrate in ACSF for 60 min at room temperature before imaging or recording. The ACSF solution contained (in mM): 126.0 NaCl, 3.0 KCl, 1.25 NaH_2_PO_4_, 1.0 MgSO_4_, 1.2 CaCl_2_, 26.0 NaHCO_3_, and 10.0 glucose.

### Live Imaging

300 µm live sections centered on the somatosensory sector of TRN (−1.4 mm from bregma) were imaged using Nikon or Zeiss upright microscopes with 2.5-5X objectives and Andor Zyla sCMOS cameras. Both epifluorescent and transmitted light (brightfield) images were obtained to characterize the topographical positions of the TRN cell types and their connections with thalamic relay nuclei.

To generate the group maps showing the average VP and POM projections to TRN (Fig. 2b), the TRN of each live slice was first outlined using either tdTomato expression driven by SOM-Cre (n=6 for POM, n=4 for VP) or bright field images (n=3 for POM, n=4 for VPM). Those outlines were then used to warp the slice images to a common reference TRN (with the bUnwarpJ plugin in ImageJ - described for the immunohistochemical analysis above), allowing precise alignment across mice for averaging. The central/edge boundaries were drawn at 20% and 80% of the medial-lateral distance across the TRN.

### Whole-Cell Recording Procedure

Brain slices (300 µm) were placed in a submersion-type recording chamber maintained at 32 ± 1°C and continuously superfused with oxygenated ACSF (above). Neurons were visualized for recording using DIC-IR optics with 40X water immersion objectives. Patch pipettes had tip resistances of 3-6 MΩ when filled with a potassium-based internal recording solution containing (in mM): 130.0 K-gluconate, 4.0 KCl, 2.0 NaCl, 10 HEPES, 0.2 EGTA, 4.0 ATP-Mg, and 0.3 GTP-Tris, 14.0 phosphocreatine-K (pH 7.25, ∼290 mOsm). During all recordings, pipette capacitances were neutralized. Series resistances (∼12-32) were compensated online (100% for current-clamp, 60-70% for voltage-clamp). Pharmacological agents (stated in the figure legends when utilized) were diluted in ACSF just before use and applied though the bathing solution. Voltages reported here were corrected for a 14 mV liquid junction potential. The reversal potential for GABA_A_ receptor-mediated responses in thalamic relay cells was -91 mV.

### Measurements of Intrinsic Physiological Properties

Resting membrane potentials were measured within 2 min of break-in. Steady-state potentials were adjusted to -74 mV or -84 mV with intracellular current to test physiological properties in tonic or burst mode, respectively. Input resistances (R_in_) and membrane time constants (τ_m_) were calculated from voltage responses (∼3 mV deflections) to small negative current injections (3-25 pA, 600 ms). For τ_m_, the voltage responses were fitted with a single exponential to the initial 50 ms of the response, omitting the first ms. R_in_ values were measured using Ohms law and input capacitances C_in_ were calculated as τ_m_ / R_in_.

Threshold (rheobase) currents were measured as the minimum injected currents required to discharge an action potential (AP; determined using 1 sec duration currents, 5 pA step increments). All other AP and after-AP properties were measured from the first AP discharged at the threshold current, but APs were only analyzed if discharged in the initial 200 ms). AP voltage thresholds were measured, and verified by visual inspection, as the potential at which the rate of rise became greater than 10 V/s. AP amplitudes and after-AP properties were measured relative to the threshold potential. AP widths were measured at half of the AP amplitude (threshold voltage to peak). Fast afterhyperpolarizations (fast AHPs) were measured as the most hyperpolarized potential immediately succeeding the AP. In tonic mode, afterdepolarizations (ADPs) were measured as the most depolarized potential within 20 ms of the AP, and slow AHPs (sAHPs) were measured as the most hyperpolarized potential within 100 ms after the ADP. In burst mode, the AHP following the burst was measured as the minimum potential within 150 ms following the burst. ADP and sAHP measurements were not considered if a second AP (or tonic APs after a burst) confounded the measurements.

Repetitive spiking properties in tonic and burst mode were measured using positive current steps (25-200 pA, 25 pA increments, 1 s duration). Spike frequency adaptation was quantified as an adaptation ratio (frequency of the last 2 APs divided by the frequency of the first 2 APs) averaged across all sweeps in which the frequency of the last 2 APs was 20-60 Hz.

To elicit offset bursts (Fig. 3, Extended Data Fig. 9 and Supplementary Information 1), the steady-state potential was adjusted to -74 mV with intracellular current, then 1 s duration negative currents were injected. Offset bursts were measured from trials in which negative current (20-300 pA, 20-25 pA test increments) led to an average voltage of -92 ± 2 mV at the end of the current step.

### Photostimulation

Synaptic physiology experiments were performed on inputs to TRN cells from excitatory thalamic relay neurons (Figs. 2,3), and on inhibitory outputs of TRN cells to thalamic relay neurons. In both cases, ChR2 was optically excited using white light-emitting diodes (LEDs) (Mightex LCS-5500-03-22) controlled by Mightex LED controllers (SLCAA02-US or BLS-1000-2). The light was collimated and reflected through a 40X water immersion objective, resulting in a spot diameter of ∼400 µm and a maximum LED power at the focal plane of 29.2 mW. The stimuli, delivered as 10 Hz trains of 1 ms flashes, were typically directed at ChR2-expressing presynaptic terminals by centering the light spot over the recorded postsynaptic cells (the postsynaptic cells did not express ChR2).

In a subset of experiments (Extended Data Fig. 6 a-c), within-cell comparisons were made of TRN cell responses to stimulation directed at opsin-expressing presynaptic axons/terminals (from VP or POM) versus stimulation directed further upstream, at or near the VP or POM cell bodies of origin. These experiments tested whether short-term synaptic plasticity differed when optogenetically stimulating soma/proximal axons of presynaptic cells versus their terminal boutons, as has been shown for some pathways.^47^

### Simulated Synaptic Current Injections

Simulated excitatory postsynaptic current (EPSC) waveforms for thalamic (VP and POM) inputs to TRN were generated from averaged measurements of optogenetically evoked synaptic responses (recorded in voltage clamp at -84 mV) from central and edge TRN cells (as in Fig. 3). These simulated synaptic currents for VP and POM inputs were matched for total synaptic charge across the 10 Hz trains (49.5 pC) and injected into the central and edge TRN cells to test features of integration in each cell type (described in the main text and legend of Fig. 4).

### Data Analyses

Analyses of electrophysiological data were performed in CED Signal 6, Molecular Devices Clampfit 10, Matlab, and Microsoft Excel. Analyses of anatomical data were performed in ImageJ (plugins used: CellCounter, ROIManager, bUnwarpJ), and Microsoft Excel.

### Statistical Analyses

Statistical comparisons were performed in GraphPad Prism7 or SigmaPlot. Statistical tests used are indicated in the Results and Figure Legends. For representation of group data, center values are means and error bars show standard error of mean (SEM). Statistical significance was defined as p <0.05, unless otherwise noted in the Results.

